# Ca^2+^-evoked sperm cessation determines embryo number in mammals

**DOI:** 10.1101/2024.09.16.613169

**Authors:** Hitomi Watanabe, Daiya Ohara, Shinichiro Chuma, Yusuke Takeuchi, Tomoatsu Takano, Ami Katanaya, Satohiro Nakao, Akihisa Kaneko, Takatoku Oida, Tatsuya Katsuno, Takuya Uehata, Toru Takeo, Munehiro Okamoto, Keiji Hirota, Gen Kondoh

**Author notes:** These authors are equally contributed (Hitomi Watanabe, Daiya Ohara, Shinichiro Chuma).

## Abstract

In mammals, fertilization takes place followed by fetus development in the maternal body, so the number of fetuses that exceeds the mother’s capacity increases the frequency of stunted growth and stillbirths, while an excessive number of pups increases postnatal stunting and maternal child-rearing stress, which is detrimental to offspring procreation and species maintenance. Therefore, various control mechanisms are thought to be at work to ensure that fertilization occurs in appropriate numbers. Sperm are not capable of fertilization immediately after they are produced in the testis, and must undergo a stepwise activation process including capacitation, hyper motility activation or acrosome reaction before they meet the egg in the oviduct, where fertilization takes place, contributing positively on fertilization, which in turn ensures the number of fetuses for the maintenance of species. In contrast, in this study, we found that a subpopulation of sperm is derived in a Ca^2+^-dependent manner during the process for sperm to acquire fertility, which may regulate the number of fetuses. When sperm were harvested from the epididymis of mice and activated *in vitro*, a subpopulation of sperm emerged from the sperm population in which Lypd4 was expressed on the sperm surface at a ratio up to 30%. The sperm in this subpopulation were rather small in size, more permeable to dyes, and had already ceased motility. We further characterized this sperm population by surface antigen screening using various monoclonal antibodies and found the expression of several proteins, such as CD55, ICOS or Ccr3, specific to this population. The emergence of this sperm population was induced at a concentration of about 4% of Ca^2+^ in body fluids and was independent of capacitation or acrosome reaction, and also apoptotic process. This subpopulation also appeared over time in sperm ejaculated into the female body, accounting for about 50% of the sperm that reached the oviduct in an hour. When this sperm subpopulation was then removed from the entire population using anti-Lypd4 antibody followed by *in utero* insemination, the fertilization rate of the oocytes collected from the oviducts doubled. Such a sperm subpopulation was also observed in macaque monkeys, and removal of this subpopulation increased the egg penetration rate of sperm, suggesting that this sperm subpopulation exists commonly in mammals and that the mother’s acceptable fetal number is adjusted by systematically sterilizing a certain number of sperm.

## Introduction

It is estimated that approximately 30% of all infertility cases are due to unexplained male infertility, the genes and molecular mechanisms responsible for which are still unknown. Our research has focused on the role of glycosylphosphatidylinositol-anchored protein (GPI-AP) in fertilization^1, 2, 3^; GPI-APs are a class of membrane-bound proteins in which proteins with various functions localize to the cell membrane surface via glycolipid anchors^4^. GPI-AP is conserved in all eukaryotes, from mono-cellular organisms, such as yeast and protozoa, to human, and genetic studies in mono-cellular organisms have shown that loss of GPI-AP is lethal. In mammals, on the other hand, more than 150 GPI-APs are known, and the functions of their protein parts, such as hydrolytic enzymes, adhesion factors, ligands, and receptors, vary widely. Loss of GPI-AP in mouse ES cells is not lethal^5^, while conditional loss of GPI-AP in various tissues is also known to impair homeostasis and the performance of their own functions, indicating that it also plays important roles in multi-cellular organisms^6, 7, 8^. GPI-APs such as CD59, Prss21, Hyal5, Prnd, and Tex101 are expressed in mouse sperm, and deletion of Tex101, among them, results in abnormal male pregnancy due to impaired sperm passage through the uterine oviduct^9^. Deletion of folate receptor 4 (FR4), a GPI-AP in the oocyte, results in female infertility due to impaired sperm-egg fusion^10^, indicating that the GPI-anchor system is essential for the establishment of fertilization in mammals. Therefore, we conceived that the sperm-specific and most highly expressed GPI-AP is important for sperm function and set out to identify the corresponding protein.

Mammalian sperm acquire various membrane modifications in the male and female reproductive organs^11, 12^. The first critical step in this physiological process is capacitation, which begins in the female reproductive tract^13^. It is well known that during capacitation, membrane cholesterol is depleted, followed by protein tyrosine phosphorylation^14, 15, 16, 17, 18^. We previously found that the innate immune factor lipocalin 2 binds to sperm membranes and promotes capacitation in a protein kinase A-dependent manner^19^. However, other mechanisms of capacitation remain to be elucidated. The acrosome reaction (AR) is another important step. The acrosome is a Golgi-derived structure located in the apical region of the sperm head. It is composed of a continuous membrane called the inner and outer acrosomal membranes^20, 21, 22, 23^. During the AR, the outer acrosomal and plasma membranes fuse to produce hybrid membrane vesicles, followed by exposure of the inner acrosomal membrane and release of the acrosomal components. AR begins when sperm migrate between the cumulus cells or bind to the Zona Pellucida (ZP)^24^. Only sperm that have completed AR can penetrate the ZP and fuse with the oocyte.

In this study, we identified the testis-specifically expressed Ly6/Plaur domain containing 4 (Lypd4). A monoclonal antibody against this protein was then generated and FACS analysis of mouse sperm revealed a Lypd4-strongly positive sperm subpopulation at a rate up to 30% during the process for sperm to acquire fertility. These sperm expressed specific proteins on their surface and were induced by physiological concentrations of Ca^2+^ ions. This was independent of capacitation or acrosome reaction, which are known to be Ca^2+^-dependent. This subpopulation accounted for about half of the sperm reaching the oviducts, and its removal from the population doubled the *in vivo* fertilization rate. In our previous studies, we have found that the sperm population is heterogeneous, and among them, sperm with altered localization of plasma membrane rafts have higher fertilization potential^3,25^. In this study, we showed for the first time that Ca^2+^ ions induced subpopulations from mouse sperm, and that this may regulate the number of fetuses. A similar subpopulation was also found to emerge in monkey sperm, suggesting that this mechanism of regulating fetal number is common in mammals.

## Results

### Identification of Lypd4

We conceived that the sperm-specific and most abundantly expressed GPI-AP is important for sperm function and decided to identify the relevant protein. Membrane protein fractions were collected from mouse epididymal sperm, and GPI-AP was specifically isolated and concentrated by PI-PLC treatment, and SDS-PAGE revealed two major bands (molecular weight 27 kDa and 35 kDa) (Supplemental Fig. 1A). Therefore, when these bands were extracted from the gel and subjected to mass spectrometry analysis, the major component of 35 kDa band was found to be carbonic anhydrase 4 (CA4) (Supplemental Fig. 1B). CA4 is expressed ubiquitously and also known to be involved in the generation of bicarbonate ions, which are necessary for sperm capacitation^26^.

On the other hand, the major component of 27 kDa band was Lypd4 (Supplemental Fig. 1C). We next examined mRNA expression of this protein in organs throughout the body and found strong testis-specific expression (Supplemental Fig. 7A). We then generated Lypd4 knock-out mice using CRISPR-Cas9 genome editing (Supplemental Fig. 2A). Sperm from these mice showed fertilization ability similar to wild-type sperm *in vitro* (Supplemental Fig. 2B), but the fertilization rate on *in utero* insemination was drastically reduced (Supplemental Fig. 2C). In natural mating, the number of sperm migrating into the oviduct decreased to about one-tenth of the wild-type sperm, and the number of pups per litter decreased (Supplemental Fig. 2D). These observations did not differ from previous studies^27^. Thus, mouse lines deficient for Lypd4 we generated here were subsequently used as controls for the Lypd4 detection system.

Next, a monoclonal antibody against Lypd4 was generated (Supplemental Fig. 3) and directly labeled with a fluorescent dye. According to previous reports, Lypd4 is a protein that localizes to the outer acrosomal membrane^27^. Therefore, immunohistochemical staining was performed using this fluorescent-labeled antibody. As a result, almost all spermatocytes or spermatids expressed Lypd4 protein in tissue sections of testis. Interestingly, the Lypd4 protein appeared to localize to intracellular vesicles (Supplemental Fig. 4B, 4D, 4F). In spermatogonia, Lypd4 was never observed, consistent with the results of single cell analysis of RNA expression or protein expression analysis by FACS (Supplemental Fig. 4F, Supplemental Fig. 7C and Supplemental Fig. 8). In epididymal sperm, on the other hand, Lypd4 protein was observed to be localized to the acrosome, consistent with previously reports^27,28^(Supplemental Fig. 5). Moreover, testicular cells and epididymal spermatozoa were treated with a cell membrane-permeable reagent and fluorescently labelled antibodies were allowed to penetrate into the cells to detect intracellular Lypd4 protein. As a result, more than 90% of testicular cells and epididymal sperm were stained, indicating that almost all of testicular and sperm potentially harbor the Lypd4 protein (Fig. 1A, 1B and Supplemental Fig. 3B). Interestingly, spermatids expressed Lypd4 protein on the cell surface, while almost none of epididymal sperm showed the surface expression, indicating that the cell surface Lypd4 is once completely shed during spermiogenesis.

**Figure 1.**
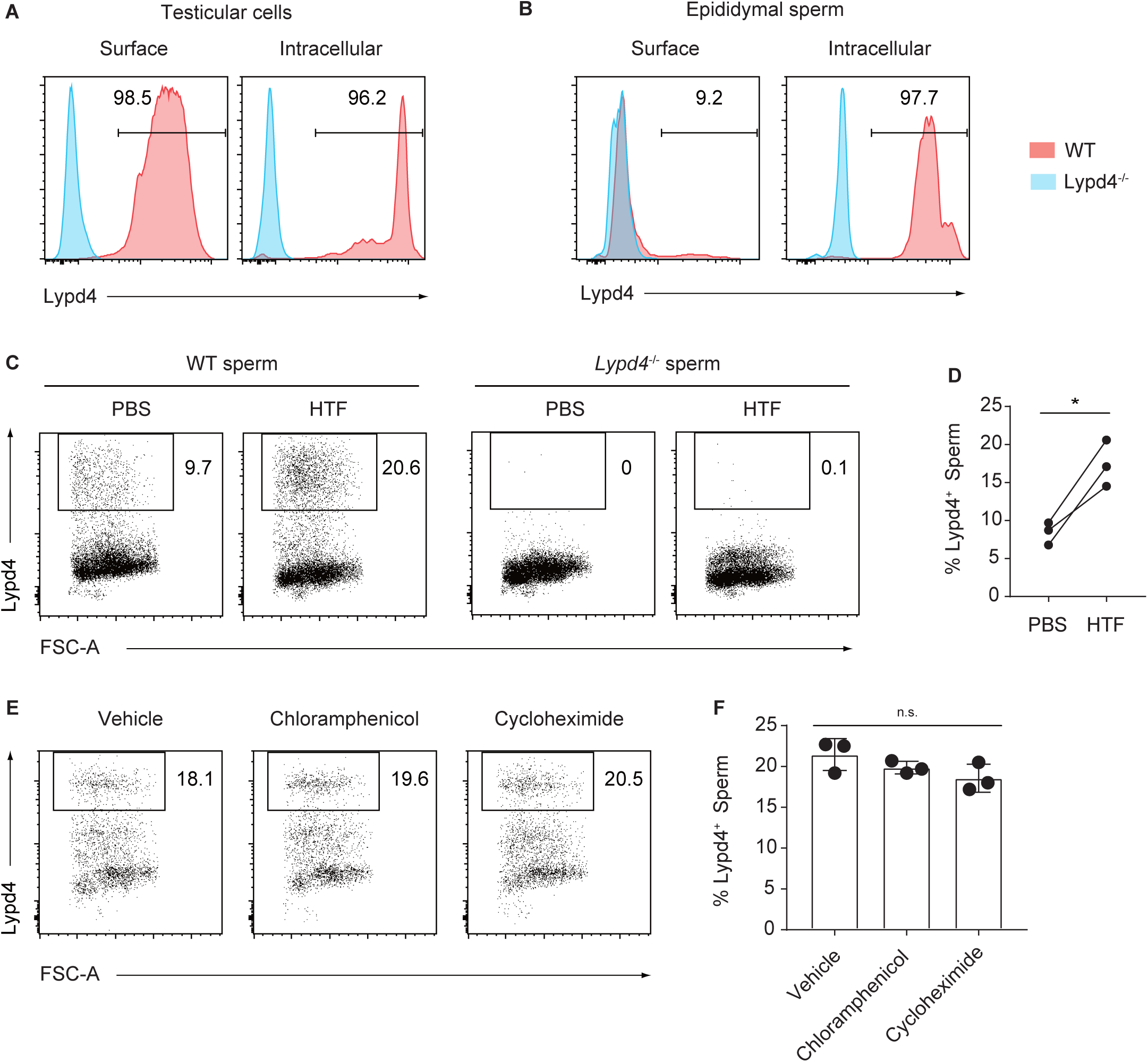
Emergence of Lypd4-positive sperm. (A) Expression of Lypd4 in testicular cells. Testicular cells showed expression of Lypd4 protein both on the cell surface and intracellularly (red shadow). Blue shadow, testicular cells of Lypd4 knock-out mice. (B) Expression of Lypd4 in epididymal sperm. Lypd4 was not observed on the sperm surface, but in almost all sperm intracellularly (red shadow). Blue shadow, epididymal sperm of Lypd4 knock-out mice. (C) In wild-type (WT), Lypd4-positive sperm certainly appear when cultured in HTF; Lypd4 knock-out sperm as a negative control. (D) There was a significant difference in the incidence of Lypd4-positive sperm between HTF (activating) and PBS (non-activating) cultures. n=3, paired t test, p=0.0308. (E) When WT sperm were cultured with chloramphenicol or cycloheximide, there was no difference in the incidence of Lypd4-positive sperm compared to the vehicle DMSO. (F) There was no difference in the incidence of Lypd4-positive sperm among DMSO only, chloramphenicol or cycloheximide treatments. n=3, one-way ANOVA followed by Tukey’s multiple comparisons test, Adjusted P value, A vs B = 0.4718, A vs C = 0.1387, B vs C = 0.5962. These results indicate that the emergence of Lypd4-positive sperm is not due to the *de novo* protein synthesis.

**Figure 2.**
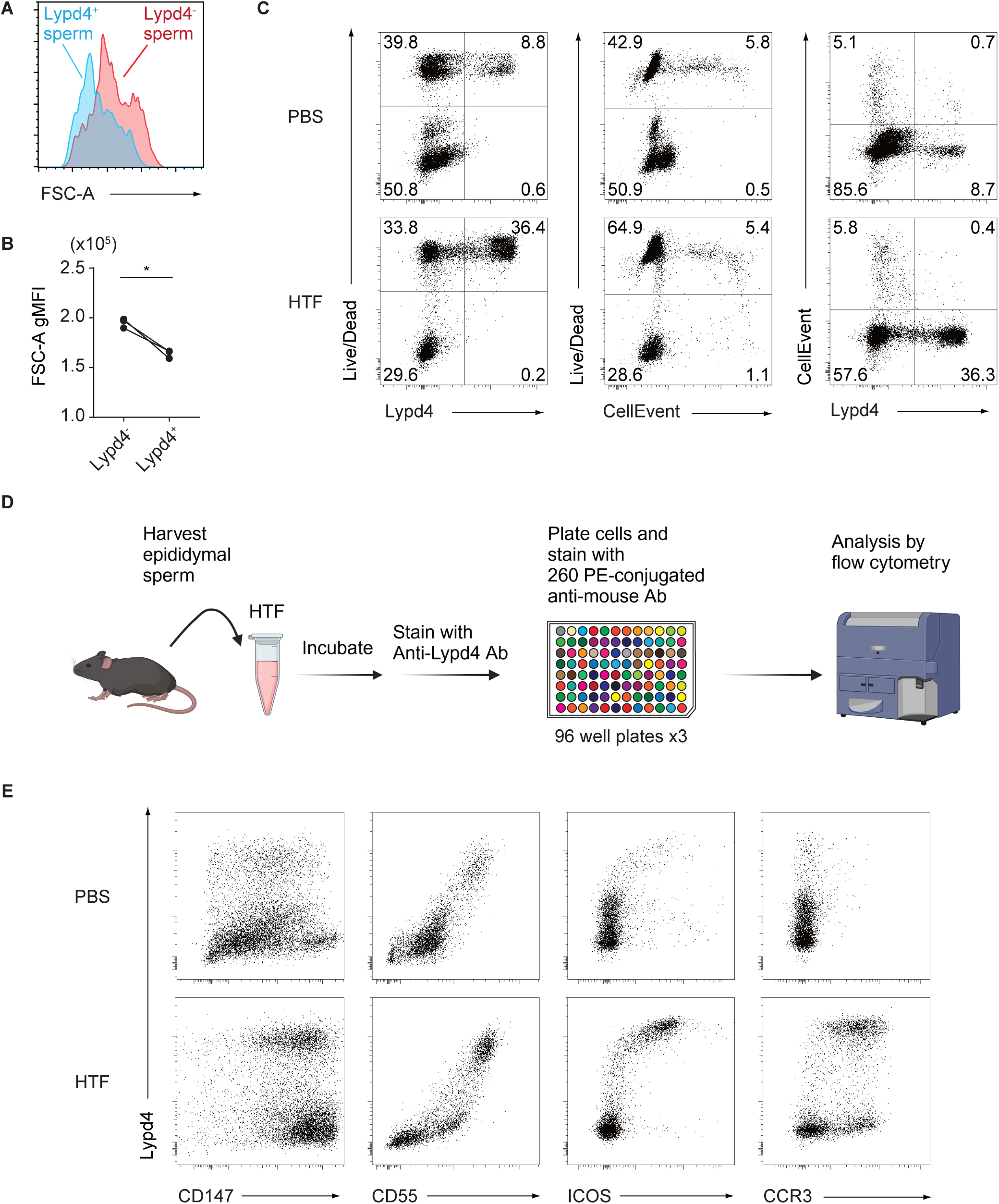
Characterization of mouse Lypd4-positive sperm. (A) Forward scattered plot (FSC) of Lypd4-positive and Lypd4-negative sperm. Lypd4-positive sperm are smaller in size. (B) Mean gMFI of FSC analysis. n=3, paired t test, p = 0.0216. (C) The wild-type sperm were cultured in HTF or PBS containing anti-Lypd4 antibody, Live/Dead reagent and CellEvent reagent, and subjected to FACS analysis. These results showed that in HTF culture, Lypd4-positive sperm were Live/Dead positive, but CellEvent negative. This indicates that Lypd4-positive sperm have enhanced membrane permeability, but cell death has not occurred. (D) FACS analysis using fluorescently labeled monoclonal antibodies against 260 different cell surface proteins. Sperm were collected from the cauda epididymis of mice and cultured in HTF medium for 2 h. Sperm were stained with anti-Lypd4 antibody and 260 independent anti-mouse antibodies, and analyzed by FACS. (E) Results of FACS analysis of CD147, CD55, ICOS, and Ccr3 are shown. The expression of CD55, ICOS and Ccr3 on the sperm surface fairly correlated with Lypd4 expression. CD147 was found to be expressed on almost all sperm surfaces by HTF activation, regardless of Lypd4 status.

**Figure 3.**
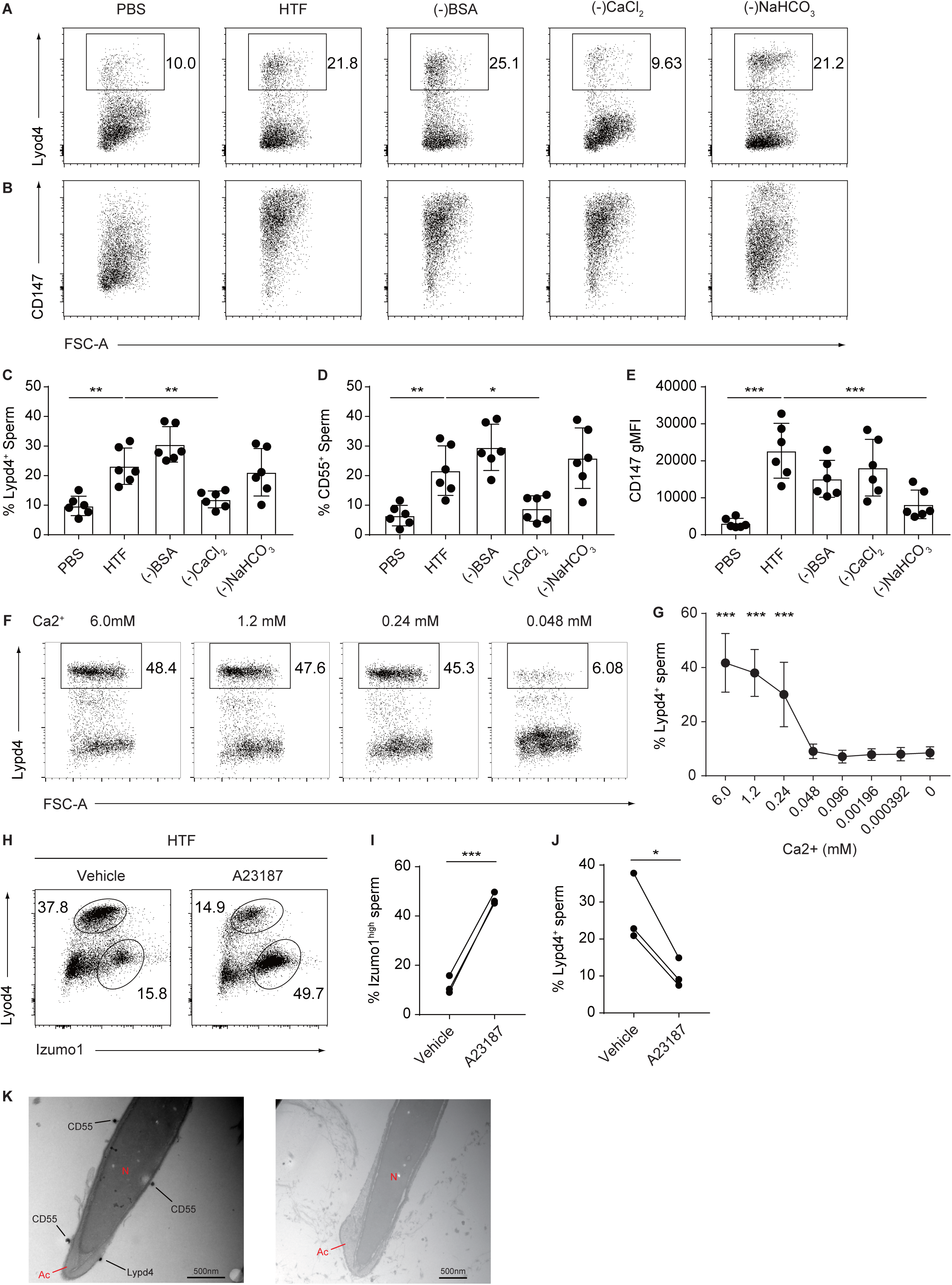
Ca^2+^ is crucial for the emergence of Lypd4-positive sperm. Each of essential components for sperm activation such as BSA, Ca^2+^, and bicarbonate was depleted from HTF, and the incidence of Lypd4-positive sperm was examined, finding Ca^2+^ is essential (A). (B) We also examined the rate of CD147-high sperm, and found that bicarbonate is an important factor for CD147-high sperm occurrence. (C) Lypd4-positive sperm incidence from multiple experiments and their statistical analysis. n=6, one-way ANOVA followed by Bonferroni’s multiple comparisons tests comparing each group with the HTF group. The results of the statistical analyses are shown in the text. (D) CD55-positive sperm incidence from multiple experiments and their statistical analysis. n=6, one-way ANOVA followed by Bonferroni’s multiple comparisons tests comparing each group with the HTF group. The results of the statistical analysis are shown in the text. (E) CD147-high sperm appearance indicated by gMFI from multiple experiments and their statistical analysis. n=6, one-way ANOVA followed by Bonferroni’s multiple comparisons tests comparing each group with the HTF group. The results of the statistical analysis are shown in the text. (F) Dose-dependency of Ca^2+^ concentration on the emergence of Lypd4-positive sperm. A Ca^2+^ concentration of at least 0.24 mM is required for the emergence of Lypd4-positive sperm. (G) The Ca^2+^ dose-dependent curve for the incidence of Lypd4-positive sperm. n=5, one-way ANOVA followed by Bonferroni’s multiple comparisons tests comparing each group with the 0mM Ca2+ group. The results of the statistical analysis are shown in the text. (H) Lypd4-positive sperm status during acrosome reaction. The progression of acrosome reaction was monitored by the expression intensity of Izumo1. In the control (Vehicle), where the acrosome reaction is slow, Izumo1 expression in Lypd4-positive spermatozoa is broadly intense. When A23187 was used to force acrosome reaction, the number of Lypd4-positive sperm decreased drastically and shifted to Izumo1 alone high. (I) Incidence of Izumo1-high sperm from multiple experiments as in (H) and their statistical analysis. n=3, paired t test, p = 0.0007. (J) Status of Lypd4-positive sperm from multiple experiments as in (H) and their statistical analysis. n=3, paired t test, p = 0.0329. (K) Immuno-electron microscopy of sperm revealed the expression of Lypd4 and CD55 on the sperm surface, where no acrosome reaction had occurred (Left). Right, Lypd4:CD55-negative sperm ongoing acrosome reaction. Ac, acrosome; N, nucleus. Scale bar, 500 nm.

**Figure 4.**
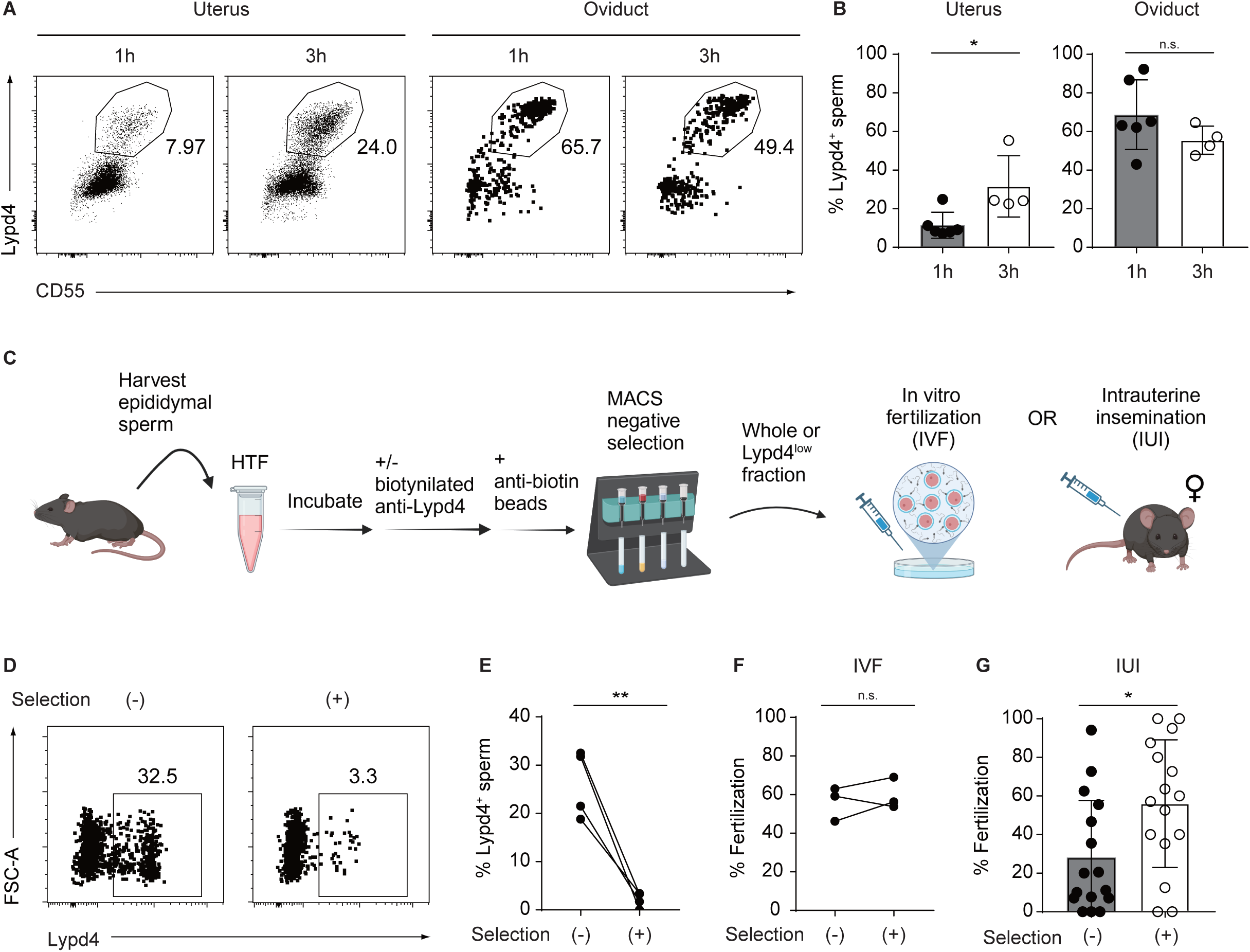
Properties of mouse Lypd4:CD55-positive sperm *in vivo*. (A) Rate of Lypd4:CD55-positive sperm in sperm ejaculated into the female body. Female mice were mated and sperm in the uterus and oviducts were collected one or three hours after mating and measured the rate of Lypd4:CD55-positive sperm by FACS. (B) Lypd4-positive sperm appearance rates from multiple experiments as in (A) and their statistical analysis. Uterus 1h n=6, 3h n=4, unpaired t test, p=0.0227; Oviduct 1h n=6, 3h n=4, unpaired t test, p=0.2074. (C) Removal of Lypd4:CD55-positive sperm by magnetic beads/magnet column. Epididymal sperm are collected and incubated at 37°C for 1 hour. Then, the appropriate amounts of biotinylated anti-Lypd4 antibody and streptavidin-labeled magnetic beads are added and incubated. Then, the antibody-treated samples are subjected to MACS MS column (Milteny). The flow through fractions were collected. After sperm counts, appropriate number of sperm were applied to IVF or IUI. (D) Using MACS column-mediated depletion, almost all Lypd4-positive sperm were removed. (E) The same experiments as in (D) was performed on four independent mice. n=4, Paired t test, p=0.0084. (F) Lypd4-positive sperm were removed and *in vitro* fertilization (IVF) was performed, but there was no significant difference in fertilization rate compared to non-removed group. n=3, Paired t test, p=0.5243. (G) Lypd4-positive sperm were removed and IUI was performed. Sperm from which Lypd4-positive sperm were removed had a two-fold higher fertilization rate with a significant difference. n=16, Mann-Whitney test p=0.0238.

### Emergence and characterization of Lypd4-positive sperm

However, when the collected sperm were treated with human tubular fluid medium (HTF), which activates sperm to be fertile, for 60 minutes, up to 20% of the sperm expressed Lypd4 protein on their surface (Fig. 1C and 1D). From now on, such sperm will be referred to as Lypd4-positive sperm. These sperm were relatively small, stained with a membrane-penetrating dye by FACS analysis (Fig. 2A and 2B), and had already ceased motility when observed under the microscope (Supplemental Video 1 and 2). In contrast, some sperm ceased by incubating in non-activating PBS, but showed no increased permeability for dyes (Fig.2C), suggesting that the emergence of Lypd4-positive sperm with increased membrane permeability is specific for sperm activating process. Since the membrane permeability of Lypd4-positive sperm considerably increased, we considered the possibility that these sperm characteristics could be due to apoptosis, but this possibility was ruled out because these sperm were not stained by apoptotic cell staining dye (Fig. 2B). We then examined the process by which Lypd4-positive sperm emerge in following studies. First, epididymal sperm were collected and activated in the presence of the protein translation inhibitors chloramphenicol or cycloheximide. As a result, Lypd4-positive sperm appeared in similar proportions as controls (Fig. 1E and 1F), indicating that Lypd4 is not newly synthesized during the process of sperm activation and does not appear on the sperm surface due to increased protein levels. We then fractionated the testicular cells using DNA content as an indicator and examined the amount of Lypd4 protein in each fraction by FACS analysis, showing that Lypd4 protein was widely expressed from pachytene stage spermatocytes to spermatids (Supplemental Fig. 7). Moreover, data-based single-cell RNA analysis of testicular cells revealed that Lypd4 mRNA expression was highly detected in spermatocytes to spermatids (Supplemental Fig. 8). The expression of mRNA levels within these cells was almost uniform, indicating that the appearance of Lypd4-positive spermatozoa in the sperm population is not due to the presence of cells expressing particularly high levels of Lypd4 mRNA in the testis. At the same time, we performed a single-cell RNA analysis on human testicular cells, finding similar expression profile of Lypd4 as well as CD55 or Izumo1, suggesting that testicular expressions of these molecules are conserved among mammals (Supplemental Fig. 9). Again, there are no cells that express particularly high levels of Lypd4, and since the Lypd4 protein is once lost from the sperm membrane at the epididymal sperm stage, suggesting that the emergence of Lypd4-positive sperm is due to the translocation of the Lypd4 protein from the intracellular region to the sperm surface during the process of sperm acquiring fertility.

To clarify the nature of these spermatozoa, we next performed FACS analysis using fluorescently labeled monoclonal antibodies against 260 different cell surface proteins. Sperm were collected from mouse epididymis, incubated in activating (HTF) and non-activating (PBS) culture media for 1 hour, and then live sperm were stained with Lypd4, followed by individual staining with 260 different antibodies and subjected to FACS (Fig. 2D). The expression of CD55, ICOS and Ccr3 on the sperm surface fairly correlated with positive conversion of Lypd4 (Fig. 2E). Among them, a GPI-anchored protein CD55 correlated quite strongly with Lypd4 (Fig. 2E). Then, the expression of CD55 in the testis was also examined by single cell RNA analyses and FACS analyses after fractionation of testicular cells by DNA content, and we found some overlap with Lypd4-expressing cells (Supplemental Fig. 6 and Supplemental Fig. 7). Thus, Lypd4-positive sperm are hereafter referred to as Lypd4:CD55-positive sperm in mouse. In addition, CD147 was found to be expressed on almost all sperm surfaces by HTF sperm activation, regardless of Lypd4 status. These results indicate that Lypd4:CD55-positive sperm form a distinct population from Lypd4:CD55-negative sperm because of their specific expressions of multiple membrane surface proteins.

We further investigated the conditions required for the appearance of Lypd4-positive sperm. First, we examined the requirement of BSA, Ca^2+^, or bicarbonate ion, which are all essential for sperm activation. By removing each from the HTF sperm activation medium, we found that Lypd4:CD55 positive conversion was still observed in the absence of BSA or bicarbonate ion, but not in the absence of Ca^2+^ (Fig. 3A, 3C and 3D). This observation was highly specific, as it was induced even at one-twenty fifth of the calcium concentration of HTF or body fluids (Fig. 3F and 3G). Since BSA and bicarbonate ion are essential for inducing capacitation, the Lypd4:CD55 positive conversion is not associated with capacitation. In contrast, membrane surface conversion of CD147, a molecule known to be important for spermatogenesis^29^, was not Ca^2+^-dependent, but apparently decreased upon removal of bicarbonate ion, suggesting association with capacitation (Fig. 3B and 3E).

Next, we examined the relationship with the acrosome reaction, which is known to be induced by forced Ca^2+^ influx into sperm. Epididymal sperm were collected and incubated in the presence of Ca^2+^ and calcium ionophores A23187, and the induction of acrosome reaction was examined using changes in Izumo1 staining as an indicator^30^. First, the status of Izumo1 was analyzed by FACS using fluoro-Izumo1 antibody during HTF treatment, showing a broad-ranged staining pattern in Lypd4:CD55-positive sperm (Fig. 3H). This may indicate that Izumo1 antibody penetrated into a certain number of sperm, which was thought to be related to the dye penetration shown in Fig. 2B. A certain amount of Lypd4-negative sperm also became positive for Izumo1, indicating that the acrosome reaction occurs spontaneously regardless of Lypd4 expression (Fig. 3H). We then induced acrosome reaction by adding A23187 to HTF and performed FACS analysis. As a result, Izumo1 expression was significantly increased, while Lypd4 expression was fairly decreased (Fig. 3I, 3J and 3K). This may be due to the fact that Izumo1, which is originally localized in the inner acrosomal membrane, is exposed on the sperm surface during acrosome reaction, while Lypd4, which is localized in the outer acrosomal membrane, is shed out^27^. Moreover, detailed analyses by immune-electron microscopy revealed that numerous spermatozoa with sperm surface Lypd4:CD55 expression seemed noraml with intact acrosome (Fig. 3K and Supplemental Fig. 6). Interestingly, Lypd4 and CD55 did not co-localize on the sperm surface at all. Although both proteins are GPI-anchored proteins and distribute to plasma membrane rafts, it may be due to the somewhat narrow window of CD55 expression in the testicular cell population compared to Lypd4 (Supplemental Fig. 7B and Supplemental Fig. 8B). Thus, this observation may be explained by difference in the timing of protein synthesis and plasma membrane distribution.

Thus, we concluded here that the Lypd4:CD55 positive conversion is Ca^2+^-dependent but independent of capacitation and acrosome reaction, and that Lypd4:CD55-positive sperm do not undergo apoptosis and do not show morphological abnormalities at electron microscopy level, being not solely dead cells.

### Lypd4:CD55-positive sperm reduces fertilization ability of sperm population *in vivo*

Based on the results from *in vitro* experiments, we next examined whether Lypd4:CD55-positive sperm also emerge in the sperm population ejaculated into the female. We co-housed wild-type male and female mice and collected sperm from the uterus and oviducts after confirming mating, stained them with fluorescently labeled anti-Lypd4 and anti-CD55 antibodies, and performed FACS analysis. Here we found that Lypd4:CD55 positive sperm appeared one hour after copulation and reached to 30% after three hours (Fig. 4A and 4B). At the same time, Lypd4:CD55-positive sperm were also appeared in the oviducts, and eventually about half of the sperm in the oviducts were Lypd4:CD55 positive (Fig. 4B). These findings suggest that Lypd4:CD55-positive sperm consist a significant subpopulation *in vivo* and reach and function in the oviducts, the site where fertilization occurs.

Then, we evaluated the fertilization potential of Lypd4:CD55-positive sperm. Here we removed Lypd4:CD55-positive sperm from the sperm population using anti-Lypd4 antibody and compared the fertilization ability with that of the non-removed sperm population. HTF-activated sperm were serially treated with biotin-conjugated anti-Lypd4 antibody and magnetic beads coupled with an anti-biotin antibody, applied to a magnetic column trapping Lypd4:CD55-positive sperm, and then the flow through fraction containing only Lypd4:CD55-negative sperm was used to evaluate fertilization ability (Fig. 4C). Lypd4:CD55-positive sperm were fairly removed from the sperm population with an efficiency of over 90% (Fig. 4D and 4E). Therefore, we first compared the *in vitro* fertilization performance of the two fractions: the flow-through fraction from which the Lypd4:CD55-positive sperm were removed and the control fraction without anti-Lypd4 antibody which they were not removed, and the fertility of both fractions were examined by incubating with ovum. As a result, there was no difference in the *in vitro* fertilization ability between two fractions (Fig. 4F).

To investigate the contribution of Lypd4:CD55-positive sperm to fertilization *in vivo*, we attempted *in utero* insemination. Biotin-conjugated anti-Lypd4 antibody was applied to HTF-activated sperm, which were then incubated with magnetic beads coupled with anti-biotin antibody, applied to a magnetic column, and then flow through fraction was administered into the uterus of estrous female mice (Fig. 4C). When a sperm population containing no Lypd4-positive sperm was administered, the fertilization rate was twice as high as that of a population containing Lypd4:CD55-positive sperm (Fig. 4G), indicating that Lypd4:CD55-positive sperm may reduce *in vivo* fertilization rate by hindering fertile sperm to access ovum.

### The sperm subpopulation appears in monkey upon sperm activation

In order to investigate whether the emergence of subpopulation associated with sperm activation is common among mammals, we further performed surface antigen screening on monkey sperm. Here, we used sperm of Japanese macaque monkeys, which are genetically far from mice but more closely related to humans. Macaque monkey proteins are generally homologous to human proteins, so anti-human antibodies can be used. First, we generated four independent anti-human LYPD4 monoclonal antibodies that may detect macaque LYPD4. Although LYPD4 protein was detected in the monkey testis, HTF-activated monkey sperm did not express LYPD4 at all, different from mouse sperm (Supplemental Fig. 10).

Then, we examined expression of CD55, which was found to be strongly correlated with Lypd4 expression in mouse sperm. Epididymal sperm were collected from anesthetized monkeys and treated with HTF medium for 2h to acquire fertilization ability. Sperm expressing CD55 protein on the sperm surface appeared in 60∼80% (Fig. 5A and 5B). From now on, we will refer to these sperm as CD55-positive sperm.

**Figure 5.**
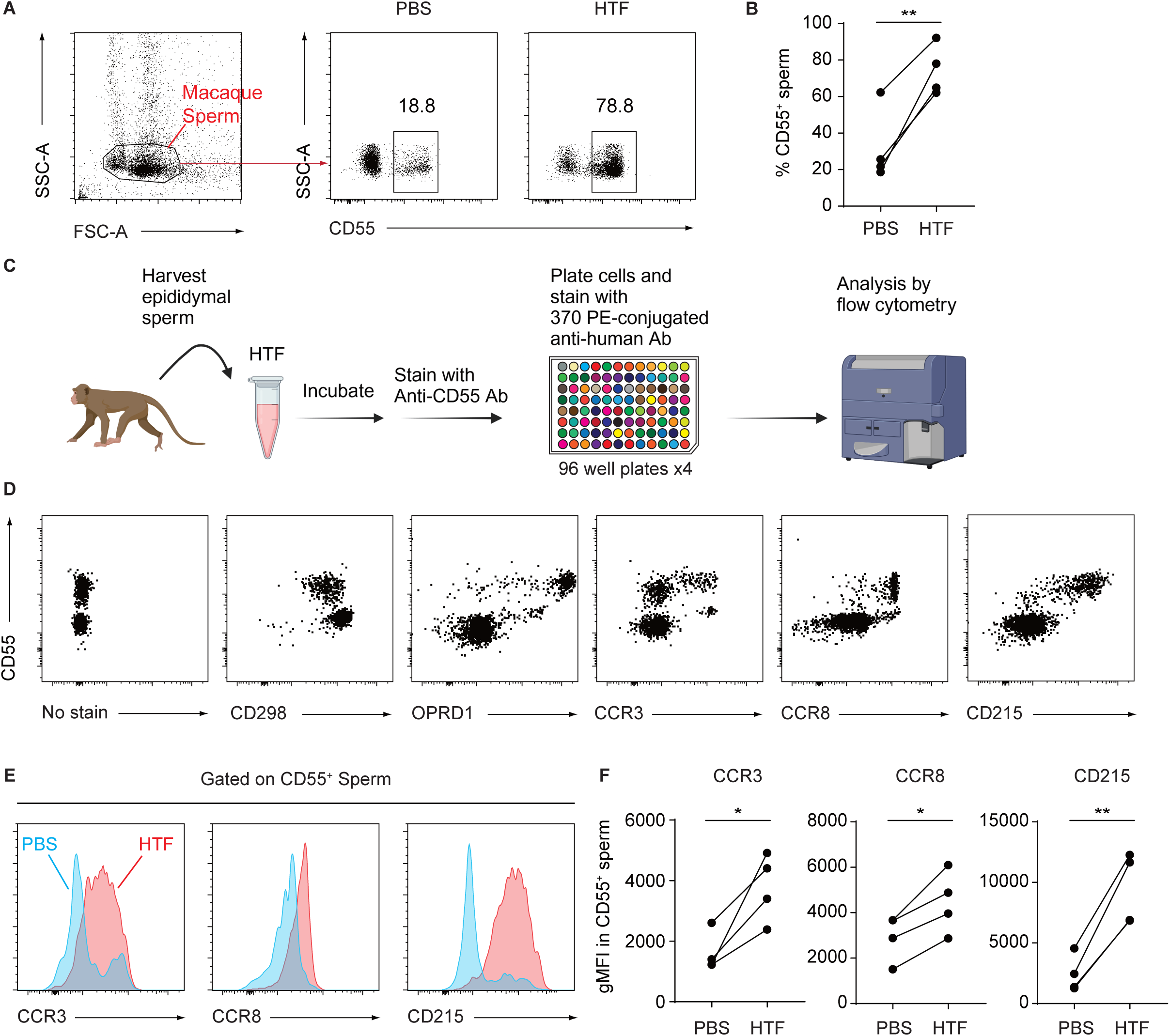
Properties of monkey CD55-positive sperm. (A) Sperm were collected from monkey epididymis under anesthesia and cultured in HTF. As there was no expression of Lypd4 in monkey sperm (Supplemental Fig. 10), it was examined with CD55. After two-dimensional expansion with side scattered (SSC) and forward scattered (FSC) as parameters, monkey sperm fractions were taken and examined for CD55 expression on the sperm surface, which showed a drastic increase in the incidence of CD55-positive sperm. (B) The same analysis as in (A) was performed on four independent monkeys. There was a significant difference in the incidence of CD55-positive sperm between HTF (activating) and PBS (non-activating) cultures. n=4, paired t test, p=0.0069. (C) FACS analysis using fluorescently labeled monoclonal antibodies against 370 different cell surface proteins. Sperm were collected from the epididymis of monkey and cultured in HTF medium for 2 h. Sperm were stained with anti-CD55 antibody and 370 independent anti-human antibodies, and analyzed by FACS. (D) Results of FACS analyses of CD298, OPRD1, CCR3, CCR8, and CD215 (IL15Rα) expressions along with CD55. The expression of OPRD1, CCR3, CCR8, and CD215 on the sperm surface fairly correlated with membrane translocation of CD55. CD298 was found to be expressed on almost all sperm surfaces by HTF, regardless of CD55 status. (E) Sperm were collected from monkey epididymis under anesthesia and cultured in HTF or PBS. The CD55-positive sperm fractions were then examined for the expression of CCR3, CCR8 and CD215, finding that the incidence of positive sperm was higher in HTF than in PBS in all these proteins. (F) The same analysis as in (E) was performed on four independent monkeys. There was a significant difference in the incidence of CCR3-, CCR8-, or CD215-positive sperm between HTF (activating) and PBS (non-activating) cultures. n=4 each, paired t test CCR3 p= 0.0281, CCR8 p=0.0156, CD215, p=0.0043.

Next, we performed FACS analysis using fluorescent-labeled monoclonal antibodies against 370 different cell surface antigens to clarify the difference between CD55-positive and CD55-negative sperm as in mouse. Sperm were collected from monkey epididymis, incubated in activating (HTF) and non-activating (PBS) culture medium for 2h, and then live sperm were stained with CD55 and individually stained with 370 different antibodies for FACS analysis (Fig. 5C). We found expressions of OPRD1, CCR3, CCR8 and CD215 (IL15 receptorα) correlated with CD55 on the sperm surface (Fig. 5D). CD55-positive sperm fractions were then examined for expression of CCR3, CCR8 and CD215, and the incidence of positive sperm was higher in HTF than in PBS for all these proteins (Figures 5E and 5F), indicating that CD55-positive sperm are alive, although their motility is arrested.

These results indicate that monkey CD55-positive sperm are also distinct from CD55-negative sperm, like mouse Lypd4:CD55-positive sperm.

### CD55-positive sperm reduces fertilization ability of sperm population in monkey

We next evaluated the fertilization potential of CD55-positive sperm. As in the mouse experiment, we prepared a mixture of phycoerythrin (PE)-cy7-conjugated anti-CD55, PE-conjugated anti-OPRD1, PE-conjugated anti-CCR3, PE-conjugated anti-CCR8 and PE-conjugated anti-CD215 (IL15Rα) antibodies for removing CD55-positive sperm subpopulation. Sperm were incubated with these antibodies followed by contact with magnetic beads conjugated with anti-PE antibody, and then applied to a magnetic column to trap CD55-positive sperm and collect flow through fractions (Fig. 6A). Here, CD55-positive sperm were removed with an efficiency of more than 90% (Fig. 6B and 6C). Sperm populations containing CD55-positive sperm and not containing CD55-positive sperm were then contacted with zona pellucida removed hamster ovum to measure sperm penetration ability. This hamster test is known to be the most reliable parameter of human sperm fertilization ability in clinical practice^31^. Finally, the administration of a sperm population that did not contain CD55-positive sperm subpopulation resulted in a two-fold higher sperm penetration rate than a population containing them (Fig. 6D and 6E). Since CD55-positive sperm had also ceased motility when observed under the microscope (Supplemental Video 3 and Supplemental Video 4), monkey CD55-positive sperm is also an obstacle for fertile sperm to access ovum, like in mouse.

**Figure 6.**
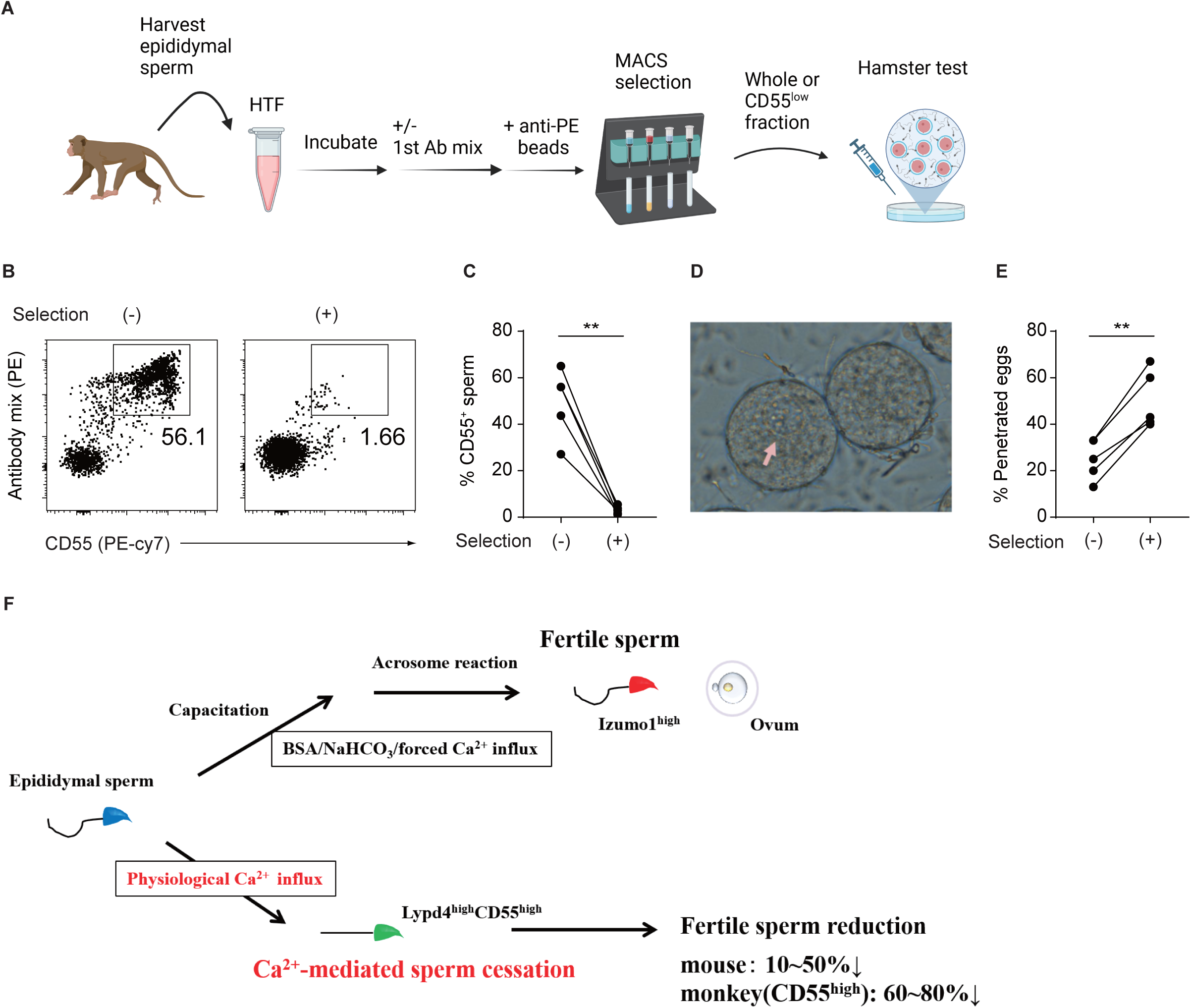
CD55-positive sperm reduces fertilization ability of sperm population in monkey. (A) Removal of CD55-positive sperm by magnetic beads/magnet column. Epididymal sperm are collected incubated at 37°C for 1 hour. Then, the appropriate amounts of PE-cy7 conjugated anti-CD55, PE-conjugated anti-OPRD1, PE-conjugated anti-CCR3, PE-conjugated anti-CCR8 and PE-conjugated anti-CD215 antibodies coupled with anti-PE conjugated magnetic beads, are added and incubated. Then, the antibody-treated samples are subjected to MACS MS column. The flow through fractions were collected. After sperm counts, appropriate number of sperm were applied for the hamster test. (B) Using MACS column-mediated depletion, almost all CD55-positive sperm were removed. (C) The same experiments as in (B) was performed on five independent monkeys. n=5, paired t test, p=0.0019. (D) Representative hamster egg after monkey sperm penetration. The pronucleus with numerous nucleoli can be seen (arrow). (E) CD55-positive sperm were removed and the hamster test was performed. Sperm from five independent monkeys from which CD55-positive sperm were removed showed a twofold higher penetration rate with a significant difference. n=5, paired t test p=0.0010. (F) Summary schematic of this study. Epididymal sperm that have not yet been activated are negative for Lypd4:CD55 on the sperm surface, which is due to their being shed during spermiogenesis. When these sperm were ejaculated into female reproductive tract, some of them readily arrest in the presence of physiological concentrations of Ca^2+^ ion. In this sperm subpopulation, Lypd4 and CD55 are translocated to the sperm surface, regardless of capacitation or acrosome reaction. Reduction in the number of fertile sperm by roughly 50% in mice and 60-80% in monkeys is thought to adjust fertilization to a particular number. On the other hand, sperm that do not cease motility undergo a conventional sperm activation process using Ca^2+^ that eventually exposes Izumo1, which is essential for fusion with the egg, on the sperm surface and finally complete fertilization.

## Discussion

### Negative regulation of fertilization by a sperm subpopulation

Many previous literatures have explained that the intracellular concentration and/or forced influx of Ca^2+^is important for inducing sperm capacitation, hyper motility activation or acrosome reaction^14, 15, 16, 17, 18, 32, 33, 34^. In contrast, here we found that Ca^2+^-managed sperm cessation occurs during sperm activation process. By summarizing our results in Fig. 6E, we display a mechanism negatively regulating fertilization by a sperm subpopulation in mammals. Epididymal sperm that have not yet been activated are negative for Lypd4 and CD55 on the sperm surface, which is due to their being shed during spermiogenesis. When these sperm were ejaculated into female reproductive tract, some of them readily arrest in the presence of physiological concentrations of Ca^2+^ ion. In this sperm subpopulation, the sperm membrane becomes fluidized, accompanied by increased membrane permeability for small molecules and translocation of Lypd4:CD55 to the sperm surface. Reduction in the number of fertile sperm by approximately 50% in mice and 60-80% in monkeys is thought to adjust fertilization to a particular number. On the other hand, sperm that do not cease motility undergo a conventional sperm activation process also using Ca^2+^ that eventually exposes Izumo1, which is essential for fusion with the egg, on the sperm surface and finally complete fertilization. In this study, we conducted experiments using protein synthesis inhibitors, FACS analysis of fractionated testicular cells, and single cell RNA analysis using publicly available data on testicular cells to elucidate the mechanism how Lypd4:CD55-positive sperm emerge, while we are unable to understand now. Further investigations are expected.

### Molecular mechanism of membrane surface translocation

Lypd4:CD55-positive spermatozoa are somewhat smaller and stained with membrane-permeable molecules, suggesting that the sperm membrane of such sperm is more fluid and undergo structural changes. Previous studies have shown that Lypd4 are localized in the outer acrosomal membrane^27^. In the present study, it is likely that Lypd4 and also CD55, which were translocated to the sperm membrane surface prior to capacitation or acrosome reaction, might be transported from the outer acrosomal membrane to the surface membrane by some Ca^2+^-dependent mechanism. As seen in the electron microscopy of Fig. 3K and Supplementary Fig. 6, the sperm membrane and acrosomal membrane are in close back-to-back contact and seem easily to exchange molecules between both membranes. Since sperm membrane and outer acrosomal membrane fuse and desorb upon acrosome reaction^35,36^, these membranes might be potentially fluid, thus suggesting Lypd4:CD55 membrane translocation exploiting this sperm membrane fluidity.

### Difference between *in vitro* and *in vivo* fertilization rates on removing Lypd4:CD55-positive sperm in mice

When Lypd4:CD55-positive sperm were removed from the mouse sperm population, there was no difference in *in vitro* fertilization (IVF) rate compared to non-removing control, but there was a difference *in vivo*. In contrast, removal of CD55-positive sperm increased the *in vitro* sperm fertility assessment in monkeys. This is probably due to the low incidence of Lypd4:CD55-positive sperm, at most 10%-30%, on sperm activation *in vitro*, where the effect of sperm removal is not fully reflected in IVF in which an excessive number of sperm is usually inseminated. On the other hand, the innate CD55-positive sperm incidence was as high as 60-80% in monkeys, suggesting such a significant effect *in vitro*. Another possibility is that Lypd4:CD55-positive sperm not only stop the motility and become sterile, but also express particular molecules that may actively suppress the fertilization ability of Lypd4:CD55-negative sperm *in vivo*. In fact, since Lypd4:CD55-positive sperm express not only Lypd4 and CD55 but also ICOS and Ccr3, a thorough analysis of these molecules may reveal such inhibitory mechanisms.

### Lypd4 is not essential for fertilization

Since Lypd4 KO sperm have shown normal *in vitro* fertilization ability and pregnancy ability is not completely lost (Supplemental Fig. 2), Lypd4 itself is not essential for fertilization. Therefore, Lypd4 is just an indicator of such sperm subpopulation whose membrane surface molecules have changed their localization in a Ca2+-dependent manner during sperm activation. In this study, multiple molecules, such as CD55, ICOS or Ccr3, that were localized on the membrane surface specifically in Lypd4-positive sperm were found. Thus, it is interesting to investigate whether they are crucial for sperm fertilization ability by generating and analyzing KO mice for each of these molecules.

### Difference in Lypd4:CD55-positive sperm emergency between mice and monkeys

As shown in Supplemental Fig. 8 and Supplemental Fig. 9, the mRNA expression profiles of Lypd4, CD55 or Izumo1 in the testis are quite similar in mouse and human, suggesting that this is conserved among mammals. However, not only do monkey sperm not express LYPD4 protein at all (Supplemental Fig. 10), but the occurrence of monkey CD55-positive sperm is roughly twice as high as mouse Lypd4:CD55-positive sperm, indicating that post-translational regulation of these genes is apparently different between species. Since Mice are prolific animals, and monkeys are small-bearing animals, it seems that the incidence of positive sperm shows inverse correlation to pup number of these animals, contributing to regulate the number of off-spring born, along with the number of ovulations per cycle. To confirm this notion, relationships between the rate of occurrence of CD55-positive sperm and the number of off-spring born awaits study in other animal species.

### Differences in membrane proteins specific to Lypd4:CD55-positive sperm in mice and monkeys

In mouse Lypd4:CD55-positive sperm, ICOS and Ccr3 were specifically membrane-converted, whereas in monkey CD55-positive sperm, OPRD1, CCR3, CCR8 and CD215 (IL15Rα) were specific, with only CD55 and Ccr3 common between mouse and monkey. These differences, as well as the differences in the presence or absence of Lypd4, may indeed speak to species differences, but on the other hand, they may also be due to the specificity and sensitivity of the antibodies used in the screening. Therefore, when using KO mice to study the function of these molecules, mouse homologues of monkey sperm-specific molecules should also be investigated.

### Application to diagnosis and treatment of human infertility

The results of this study could lead to the development of diagnostic and therapeutic methods for human infertility. In particular, antibodies against some surface antigens identified in monkey sperm could be immediately applicable to human sperm and worth testing in cases other than male infertility due to reduced sperm count or malformed sperm. The selection of sperm by specific surface markers may also increase fertilization rates in *in utero* insemination and *in vitro* fertilization. If the relationship between specific markers and fertilization rates is clarified, they could be used as diagnostic markers for infertility. Furthermore, as the development of male infertility is not considered to be a simple matter of a single molecular abnormality, mass screening using multiple antibodies, as in this study, could be useful in the development of diagnostic and therapeutic methods.

## Materials and Methods

### Animal experiments approval

All animal experiments were approved by the Animal Experiment Committees of the Institute for Life and Medical Sciences (LIME), Center for the Evolutionary Origins of Human Behavior (EHUB) and Kyoto University.

### Identification of Lypd4

Mouse epididymal spermatozoa are collected and dissolved in 50mM Tris pH 8.0, 150mM NaCl. Samples are centrifuged at 100,000×g and the precipitates are collected. Then, the precipitates are dissolved in 50mM Tris pH8.0, 1% Triton-X114, warmed to 37°C for 10 min and phase-separated. Centrifuge at 20,000×g for 10 min at 25°C, discard the water-soluble fraction in the upper layer, and collect the detergent-soluble fraction in the lower layer. An equal volume of 50 mM Tris pH 8.0, 150 mM NaCl solution is added, warmed at 37°C for 10 min to separate the phases., and then centrifuged at 20,000×g at 25°C. These treatments are repeated for three times. Finally, an equal volume of 50 mM Tris pH 8.0, 150 mM NaCl solution containing phosphatidylinositol-specific phospholipase C (PI-PLC, Molecular probe) is added and incubated at 37°C for 16 hours. After centrifugation at 20,000×g for 10 minutes at 25°C, the upper water-soluble fraction is collected. This aqueous fraction is applied to SDS-PAGE, stained with CBB, and the bands are cut out and collected (Supplemental Fig. 1). Next, in-gel trypsin digestion is performed, and the resulting peptides are collected and applied to LC-MS/MS for protein identification.

### Analysis of Lypd4 mRNA expression

RNAs were extracted from mouse tissues and cDNA were synthesized from them by reverse transcription. Quantitative PCR were then performed using the Lypd4-specific Taqman probe (Thermo Fisher).

### Generation and analysis of Lypd4-knockout mice

The Lypd4 cDNA sequence was applied to the CRISPRDirect (<u>CRISPRdirect (dbcls.jp)</u>)search site to find the target sequence for the guide RNA of Crispr/Cas9 genome editing procedure. Then, guide RNAs for the two target sequences such as Region 1: TCCTACCTCGTGTCTCCGCC and Region 2: ACAGCCGCGTCTGCAGGTCC were synthesized (FASMAC) and microinjected into fertilized eggs of C57BL/6N mice (Nihon SLC) with tracer RNA (FASMAC) and Cas9 protein (Thermo Fisher), respectively. As a result, three founder mice were obtained and three Lypd4-knockout mouse lines were established (KO1, KO2 for Region 1 and KO3 for Region 2). In this study, KO3 line was particularly used.

### *In vitro* fertilization

Epididymal sperm are collected from male mice of each genotype and incubated in HTF for 90 minutes. Eggs are collected from super-ovulated female mice and fertilized with 48,000 sperm in 200μL HTF; after 16 hours of incubation, the number of unfertilized eggs and 2-cell stage embryos are measured and the fertilization rate (% fertility=No. 2-cell embryos/ No. 2-cell emmbryos+unfertilized eggs ×100) are calculated.

### *In utero* insemination

Epididymal sperm are collected from male mice of each genotype and incubated in HTF for 30 minutes. A shot of 48,000 sperm are injected into the uterus of each super-ovulated female mouse, and eggs are harvested from the ampulla of oviduct 16 hours later. The number of unfertilized eggs and 2-cell stage embryos are measured and the fertilization rate (% fertility=No. 2-cell embryos/ No. 2-cell emmbryos+unfertilized eggs ×100) are calculated.

### Measuring litter size in natural mating

Male mice of each genotype are co-housed with C57BL/6N female mice, and those females that are found to be plug-positive are kept alone, and litter size is measured.

### Sperm counting in uterus and oviduct

Male mice of each genotype are co-housed with C57BL/6N female mice, and uterus and oviducts are harvested at indicated time after plug confirmation. Sperm were flushed out from uterine body or oviduct with PBS, stained with antibodies and then applied to FACS analyses.

### Generation of anti-Lypd4 monoclonal antibodies

Ba/F3 cells expressing mouse or human Lypd4 are inoculated multiple times into C57BL/6 mice and collect the serum. This is used to immuno-stain sperm, Ba/F3 cells, and Ba/F3 cells expressing Lypd4 (Ba/F3 cells-Lypd4) to confirm that immunity is established (Supplemental Fig. 3A). Splenocytes are harvested from immunized mice and fused with SP2/0 myeloma cells to generate hybridomas. Next, the hybridoma is diluted to the single-cell level, and after sufficient growth, the culture supernatant is harvested and screened by immunostaining for Ba/F3-Lypd4 cells. As a result, 10 clones of anti-mouse antibodies and 4 clones of anti-human antibodies were obtained (Supplemental Fig. 3B and Supplemental Fig. 10A). These clones were cultured in large quantities and purified on Protein G columns (Thermo Fisher), some of which were fluorescently labeled with Alexa-488 dye (Thermo Fisher). Sperm crushed with high-speed centrifugation were immune-stained with the purified antibodies (Supplemental Fig. 3B).

### Immunohistochemical staining

Each tissue is harvested, fixed in 2% paraformaldehyde/PBS, dehydrated in 15% and 30% sucrose/PBS, immersed in OCT compound, and flash frozen. Frozen sections are prepared and immune-stained with Alexa-488-conjugated antibodies. At the same time, nuclear staining is performed with Hoechst 33342/PBS to identify the location of sperm.

### Western blotting

Each tissue and cell are lysed in 2% SDS/PBS, centrifuged, and supernatants were collected. After SDS-PAGE of the solution containing 20μg of protein from each sample and transfer to nitrocellulose membrane, react with various primary antibodies, followed by reaction with peroxidase-labeled anti-mouse Ig antibody, and then add peroxidase substrate solution to detect expression of the protein of interest (GE Healthcare Amersham).

### Preparation of testicular cells

Mouse testes were isolated and placed in Hank’s Balanced Salt Solution (HBSS; Nakalai Tesque) for dissection. The tunica albuginea was removed, and seminiferous tubules were separated using No. 5 forceps. The isolated tubules were transferred to 5 ml of dissociation buffer (Sigma) and fragmented into approximately 1 mm pieces using fine scissors. Tubule fragments were subjected to vigorous pipetting (approx. 10 strokes) using a disposable pipette. The resulting cell suspension was mixed with 10 ml of 2% fetal bovine serum (FBS) / HBSS and filtered through a 70 μm cell strainer. The filtrate was centrifuged at 600 x g for 5 minutes at 4°C. The supernatant was discarded, and the cell pellet was resuspended in 1 ml of 2% FBS / HBSS for subsequent use.

Two-dimensional fractionations of testicular cells were performed according to the methods reported previously^37, 38^.

### Mouse sperm preparation, culture and FACS analysis

Collect two epididymal sperm in 100 μl HTF or PBS containing 0.5 mg/ml glucose (PBS) and incubate at 37°C for 1 hour. When detecting intracellular molecules, sperm membranes are permeabilized with eBioscience™ Foxp3 / Transcription Factor Staining Buffer Set (Thermo Fisher), according to the manufactural protocols. Then, add appropriate amounts of various antibodies, Live/Dead reagent solution (Thermo Fisher) and CellEvent caspase 3/7 detection reagent (Thermo Fisher), and incubate for 30 minutes. Then, add 100 μl of sperm fluid to 10 ml of PBS + 0.2% BSA warmed at 37°C and incubate for 10 minutes. Then, centrifuge at 800 rpm/10°C/5 min, remove an appropriate amount of supernatant, and perform FACS analysis using CantoII (BD biosciences), LSR Fortessa (BD biosciences), Cytoflex S (Beckman Coulter). Data analysis and output of FACS plot charts were performed by Flowjo (Tree Star Inc).

### Electron microscopy analysis

Collect two epididymal sperm in 100 μl HTF and incubate at 37°C for 1 hour. Then, appropriate amounts of anti-Lypd4 MoAb (mouse antibody), biotinylated anti-CD55 MoAb (rat antibody), and gold colloid-labeled secondary antibody for detecting Lypd4 (gold particle size 12 nm, Jackson ImmunoResearch) and gold colloid-labeled streptavidin for detecting CD55 (gold particle size 20 nm, Cytodiagnostics) were added, and incubated for 30 minutes. Then, add 100 μl of sperm fluid to 10 ml of PBS + 0.2% BSA warmed at 37°C and incubate for 10 minutes. Then, centrifuge at 800 rpm/10°C/5 min, remove an appropriate amount of supernatant, embed in IP gel (GenoStaff), and submit to electron microscopic analysis.

### Single cell-RNA sequence re-analysis

Lypd4, CD55, Izumo1 and other molecules expressions during mouse or human spermatogenesis were investigated by reanalyzing scRNA-seq datasets from GSM2928339, GSM2928340 and GSM292841^40^.

### Removal of sperm population by magnetic beads/magnet column

Epididymal sperm are collected in 100 μl HTF or PBS and incubated at 37°C for 1 hour. Then, add appropriate amounts of biotinylated anti-Lypd4 antibody coupled with anti-biotin antibody conjugated magnetic beads or PE-cy7 conjugated anti-CD55, PE-conjugated anti-OPRD1, PE-conjugated anti-CCR3, PE-conjugated anti-CCR8 and PE-conjugated anti-CD215 coupled with anti-PE antibody conjugated magnetic beads, and incubate for 30 minutes. Then, add 100 μl of sperm fluid to 10 ml of PBS + 0.2% BSA warmed at 37°C and incubate for 10 minutes. Then, centrifuge at 800 rpm at 10°C for 5 min, remove an appropriate amount of supernatant, and subject to MACS MS column (Milteny).

### Culture and staining of monkey sperm

Monkeys are anesthetized and collect epididymal sperm. Incubate the sperm in 100 μl HTF or PBS at 37°C for 2 hours. Then, add appropriate amounts of various antibodies and Live/Dead stain, and incubate for 30 minutes. Then, add 100 μl of sperm fluid to 10 ml of PBS + 0.2% BSA warmed at 37°C and incubate for 10 minutes. Then, centrifuge at 800 rpm/10℃ for 5 minutes, and remove an appropriate amount of supernatant for FACS analyses.

### Surface antigen screening

Mouse sperm are incubated with Alexa488-conjugated anti-Lypd4 antibody /Live/Dead reagent and monkey sperm with Alexa488-conjugated anti-CD55 antibody/Live/Dead reagent for 30 minutes. Then, add 100 μl of sperm fluid to 10 ml of PBS + 0.2% BSA warmed at 37°C and incubate for 10 minutes. Then, centrifuge at 800 rpm at 10°C for 5 min, remove an appropriate amount of supernatant, and react with 260 biotin-conjugated antibodies for mouse and 370 PE-conjugated antibodies for monkey individually for FACS analysis (LEGENDScreen antibody panel, BioLegend), according to the method described previously^39^.

### Hamster test

Monkeys are anesthetized and epididymal sperm are collected. Incubate at 37°C for 2 hours in HTF. The sperm population expressing sperm surface CD55, OPRD1 and CD215 (IL15Rα) is then removed by various biotinylated antibody-streptavidin magnetic beads, as in mouse sperm. 10-week-old Syrian hamsters are subjected to superovulation and unfertilized eggs are collected. Then, hyaluronidase treatment is performed to remove the cumulus cells, followed by dissolving zona pellucida by acidic Tyrode’s solution. A shot of 48,000 sperm is administrated in 200 μl HTF and incubated for 16 hours. The presence or absence of pronuclei with multiple nucleoli are then observed under a phase contrast microscope.

### Data handling and statistical analyses

**Figure 1.**

The data in A, B, E, and F are representative of two independent experiments, and the data in C and D are representative of more than three independent experiments.

Statistical analyses were performed by paired t test (D) and one-way ANOVA followed by Tukey’s multiple comparisons test (F). *P < 0.05. n.s., not significant. The Graph (F) depicts mean ± SD.

(D) n=3, paired t test, p=0.0308.

(F) n=3, one-way ANOVA followed by Tukey’s multiple comparisons test, Adjusted P value, A vs B = 0.4718, A vs C = 0.1387, B vs C = 0.5962.

**Figure 2.**

The data in A, B, C, and E are representative of two independent experiments.

Statistical analyses were performed by paired t test (B). *P < 0.05.

(B) n=3, paired t test, p = 0.0216.

**Figure 3.**

The data in A, B, F, G, H, I, J, and K are representative of two independent experiments and the data in C, D, and E are pooled from two independent experiments. Statistical analyses were performed by one-way ANOVA followed by Bonferroni’s multiple comparisons tests comparing each group with the HTF group in (C-E) and with the 0mM Ca2^+^ group in (G). Statistical analyses were performed by paired t tests (I, J). *P < 0.05, **P < 0.01, ***P < 0.001. The Graphs (C, D, E, and G) depict mean ± SD. (C) n=6, one-way ANOVA followed by Bonferroni’s multiple comparisons tests comparing each group with the HTF group.

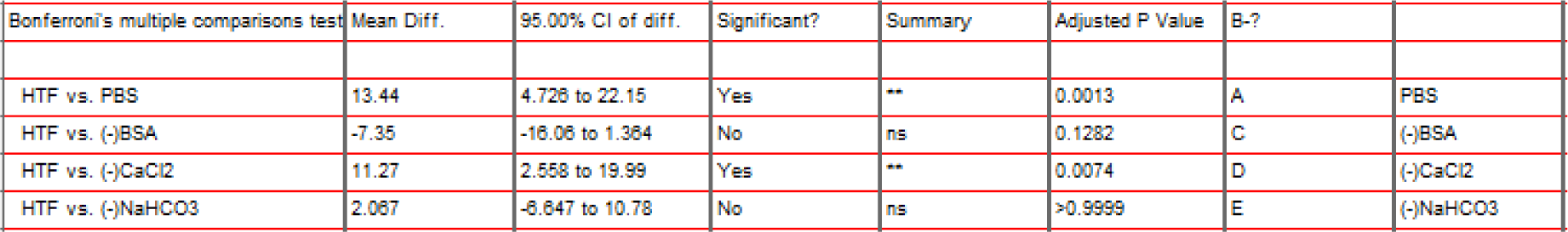

| Bonferroni's multiple comparisons test | Mean Diff. | 95.00% CI of diff. | Significant? | Summary | Adjusted P Value | B-? |  |
| --- | --- | --- | --- | --- | --- | --- | --- |
| HTF vs. PBS | 13.44 | 4.726 to 22.15 | Yes | ** | 0.0013 | A | PBS |
| HTF vs. (-)BSA | -7.35 | -16.06 to 1.364 | No | ns | 0.1282 | C | (-)BSA |
| HTF vs. (-)CaCl<sub>2</sub> | 11.27 | 2.558 to 19.99 | Yes | ** | 0.0074 | D | (-)CaCl<sub>2</sub> |
| HTF vs. (-)NaHCO<sub>3</sub> | 2.067 | -6.647 to 10.78 | No | ns | >0.9999 | E | (-)NaHCO<sub>3</sub> |

(D) n=6, one-way ANOVA followed by Bonferroni’s multiple comparisons tests comparing each group with the HTF group.

|  |  |  |  |  |  |  |  |
| --- | --- | --- | --- | --- | --- | --- | --- |
| HTF vs. PBS | 15.21 | 3.865 to 26.56 | Yes | ** | 0.0054 | A | PBS |
| HTF vs. (-)BSA | -7.867 | -19.21 to 3.48 | No | ns | 0.2953 | C | (-)BSA |
| HTF vs. (-)CaCl<sub>2</sub> | 12.82 | 1.471 to 24.17 | Yes | * | 0.0219 | D | (-)CaCl<sub>2</sub> |
| HTF vs. (-)NaHCO<sub>3</sub> | -4.233 | -15.58 to 7.114 | No | ns | >0.9999 | E | (-)NaHCO<sub>3</sub> |

(E) n=6, one-way ANOVA followed by Bonferroni’s multiple comparisons tests comparing each group with the HTF group.

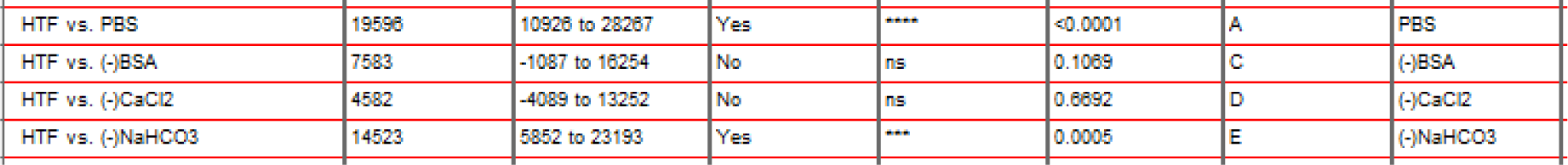

(G) n=5, one-way ANOVA followed by Bonferroni’s multiple comparisons tests comparing each group with the 0mM Ca^2+^ group.

(G) n=5, one-way ANOVA followed by Bonferroni's multiple comparisons tests comparing each group with the 0mM Ca<sup>2+</sup> group.
| Bonferroni's multiple comparisons test | Mean Diff. | 95.00% CI of diff. | Significant? | Summary | Adjusted P Value | H-? |  |
| --- | --- | --- | --- | --- | --- | --- | --- |
| 0 vs. 6.0 | -33.21 | -45.44 to -20.98 | Yes | **** | <0.0001 | A | 6.0 |
| 0 vs. 1.2 | -29.49 | -41.72 to -17.26 | Yes | **** | <0.0001 | B | 1.2 |
| 0 vs. 0.24 | -21.55 | -33.78 to -9.321 | Yes | *** | 0.0001 | C | 0.24 |
| 0 vs. 0.048 | -0.564 | -12.79 to 11.66 | No | ns | >0.9999 | D | 0.048 |
| 0 vs. 0.0096 | 1.412 | -10.82 to 13.64 | No | ns | >0.9999 | E | 0.0096 |
| 0 vs. 0.00196 | 0.648 | -11.58 to 12.88 | No | ns | >0.9999 | F | 0.00196 |
| 0 vs. 0.000392 | 0.5 | -11.73 to 12.73 | No | ns | >0.9999 | G | 0.000392 |

(I) n=3, paired t test, p = 0.0007.

(J) n=3, paired t test, p = 0.0329.

**Figure. 4.**

The data in A and D are representative of three independent experiments and the data in B, E, F, and G are pooled from more than three independent experiments. Statistical analyses were performed by unpaired t tests in (B), by paired t tests in (E and F), and by Mann-Whitney test in (G). *P < 0.05, **P < 0.01. The Graphs (B and G) depict mean ± SD.

(B) Uterus 1h n=6, 3h n=4 unpaired t test, p=0.0227. Oviduct 1h n=6, 3h n=4 unpaired t test, p=0.2074.

(E) n=4, paired t test, p=0.0084.

(F) n=3, paired t test, p=0.5243.

(G) n=16, Mann-Whitney test, p=0.0238.

**Figure. 5.**

The data in A, D, and E are representative of two independent experiments and the data in B and F are pooled from two independent experiments. Statistical analyses were performed by paired t tests in (B and F). *P < 0.05, **P < 0.01.

(B) n=4, paired t test, p=0.0069.

(F) n=4 each, paired t test, CCR3 p= 0.0281, CCR8 p=0.0156, CD215 p=0.0043.

**Figure. 6.**

The data in B and D are representative of two independent experiments and the data in C and E are pooled from two independent experiments. Statistical analyses were performed by paired t tests in (C and E). **P < 0.01.

(C) n=5, paired t test, p=0.0019.

(E) n=5, paired t test, p=0.0010.

## Acknowledgments

We thank M. Ikawa and M. Saitou for helpful discussions, and R. Takeda and M. Takenaka for extensive technical supports. This work was supported by grants from the Ministry of Education, Science, Sports, and Culture of Japan and Cooperative Research Program (Joint Usage/Research Center program) of Institute for Life and Medical Sciences, Kyoto University. Figures 2D, 4C, 5C, and 6A were created with Biorender.com.

## Author Contributions

(A) H. W., D. O., S. C., and G. K. conceived and designed experiments and wrote the manuscript. All authors performed experiments: H. W., D. O., Y. T., S. N., T. U., Tor. T. and G. K. on mouse sperm experiments; H. W., D. O., S. C., Tom. T. and Am. K. on mouse testis experiments; K. M. and Tor. T. on the mass spectrometry; H. W., D. O., Ak. K., M. O. and G. K. on monkey sperm experiments; H. W., T. O. and K. H. on monoclonal antibody generation; T. K. and G. K. on electron-microscopy. All data were analysed by D. O., S. C. and G. K.

## Supplemental Figure Legends

**Supplemental Figure 1.**
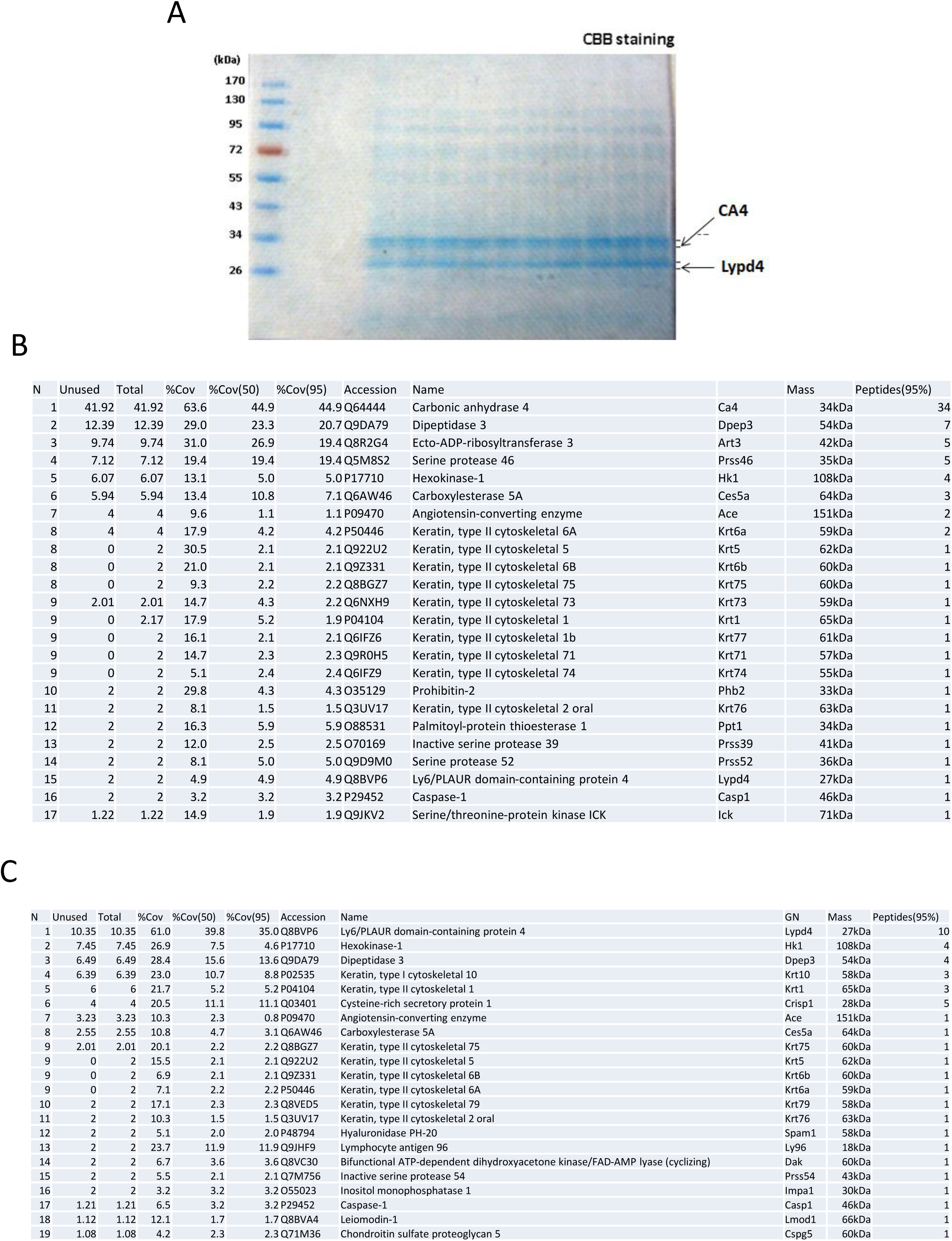
Identification of Lypd4. (A) SDS-PAGE of GPI-anchored proteins purified from epididymal sperm membrane showing major bands at 27 kDa and 35 kDa. Coomassie Brilliant Blue (CBB) staining. (B) LC-MS/MS result of the 35 kDa band. The most abundant content in this band was carbonic anhydrase 4 (CA4). (C) LC-MS/MS result of the 27 kDa band. The most abundant content in this band was Lypd4.

**Supplemental Figure 2.**
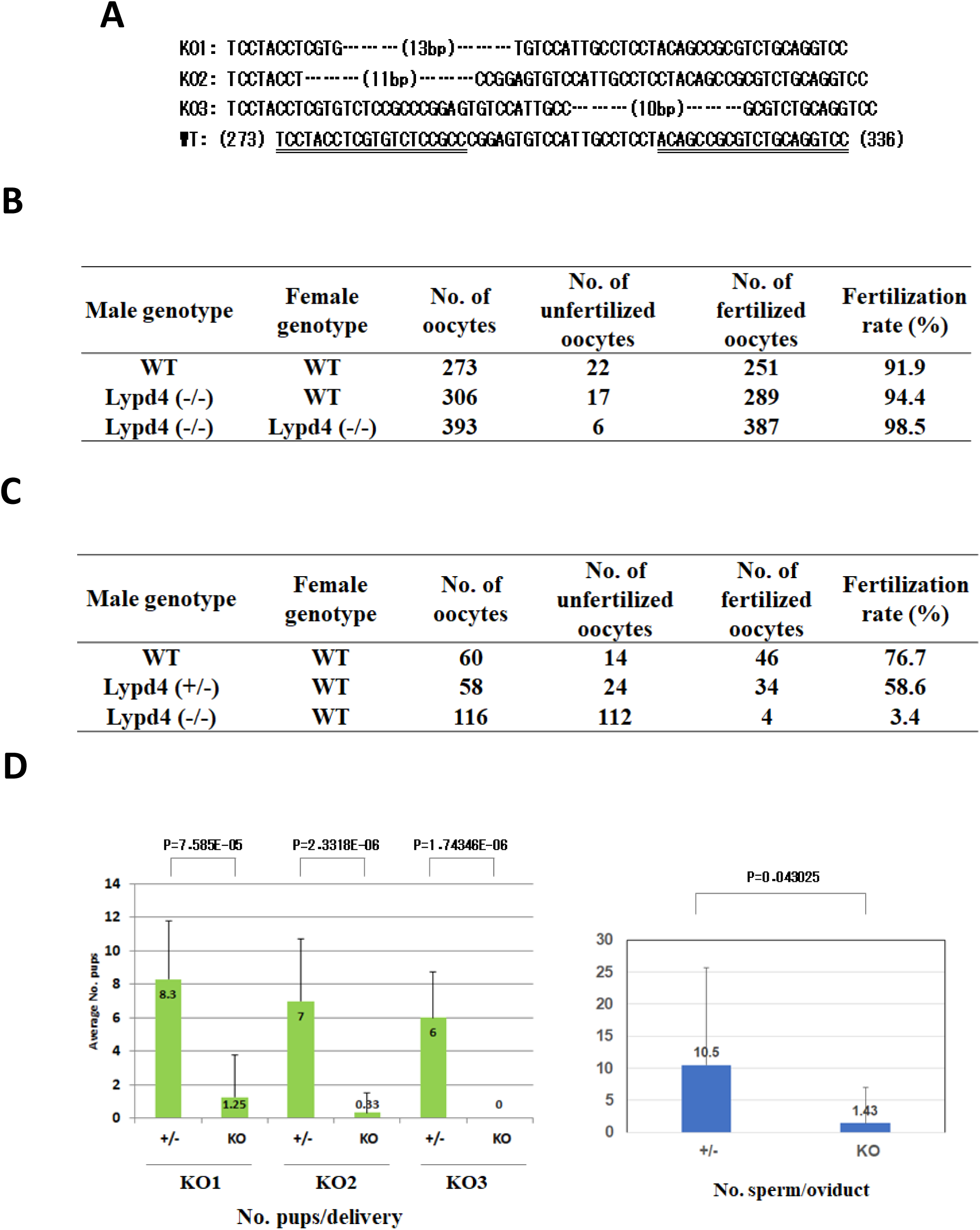
Generation and characterization of Lypd4 knock-out mouse. (A) Deletion profiles of three knock-out mouse lines (KO1, KO2, and KO3) are shown compared to wild-type (WT) DNA sequences nt. 273-336. Target DNA sequences of guide RNAs are underlined. (B) Scores of *in vitro* fertilization (IVF) using KO3 strain. Epididymal sperm from WT or Lypd4 (-/-) mice were activated with HTF and fertilized with oocytes of the respective genotypes. There were no differences in IVF rates between WT and Lypd4 (-/-). (C) Scores of *in utero* insemination using KO3 strain. Epididymal sperm from WT, Lypd4 (+/-) or Lypd4 (-/-) mice were activated with HTF and injected into the uterus of WT female mice. The next day, oocytes were collected from the ampulla of oviducts and fertilization rates were examined. Consistent with previous reports, the fertilization rate of Lypd4(-/-) sperm was significantly reduced. (D) Fertility rates in natural mating of three KO lines. Fertility was significantly lower in all KO lines compared to the respective heterozygous mice (left column each). Number of sperm collected from the oviducts of female mice mated with KO3 male mice are also shown (right). The number of sperm collected was also significantly lower than that of heterozygous mice. paired t test.

**Supplemental Figure 3.**
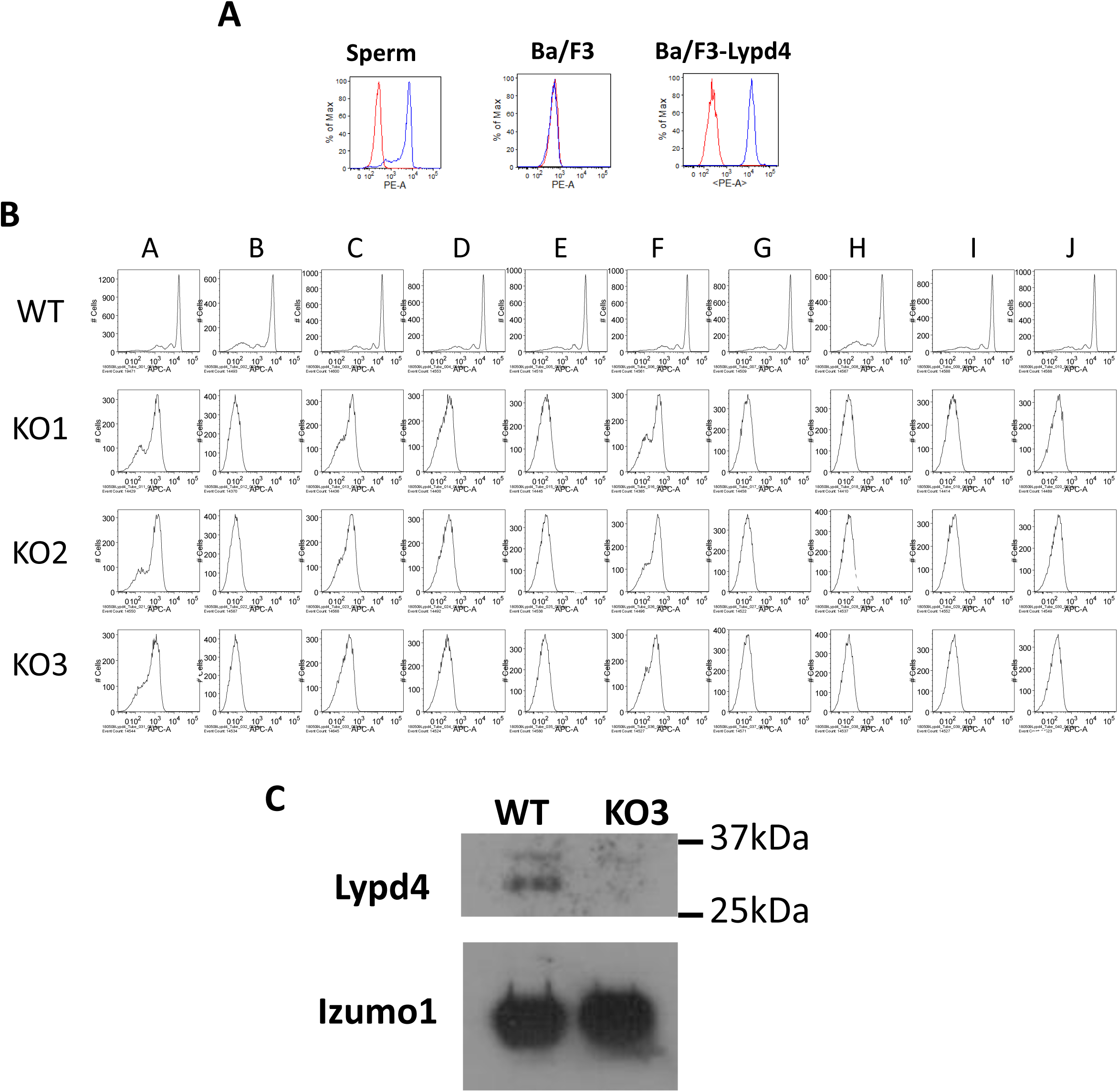
Generation of monoclonal antibodies against mouse Lypd4. (A) C57BL/6 mice immunized with Ba/F3-mLypd4 cells were successfully immunized, and serum from those mice reacted with sperm and Ba/F3-mLypd4 cells (blue line). On the other hand, they did not react with Ba/F3 cells that did not express Lypd4. Red lines indicate negative controls for experiments with pre-immune sera. (B) Ten independent clones of monoclonal antibodies were generated. These were reacted with membrane-permeabilized sperm, and all of them reacted. On the other hand, they did not react at all with sperm from three KO mouse lines. (C) Western blotting using clone J. It reacted with wild-type sperm but not with KO3-derived sperm. Quantitative controls by Izumo1 were displayed.

**Supplemental Figure 4.**
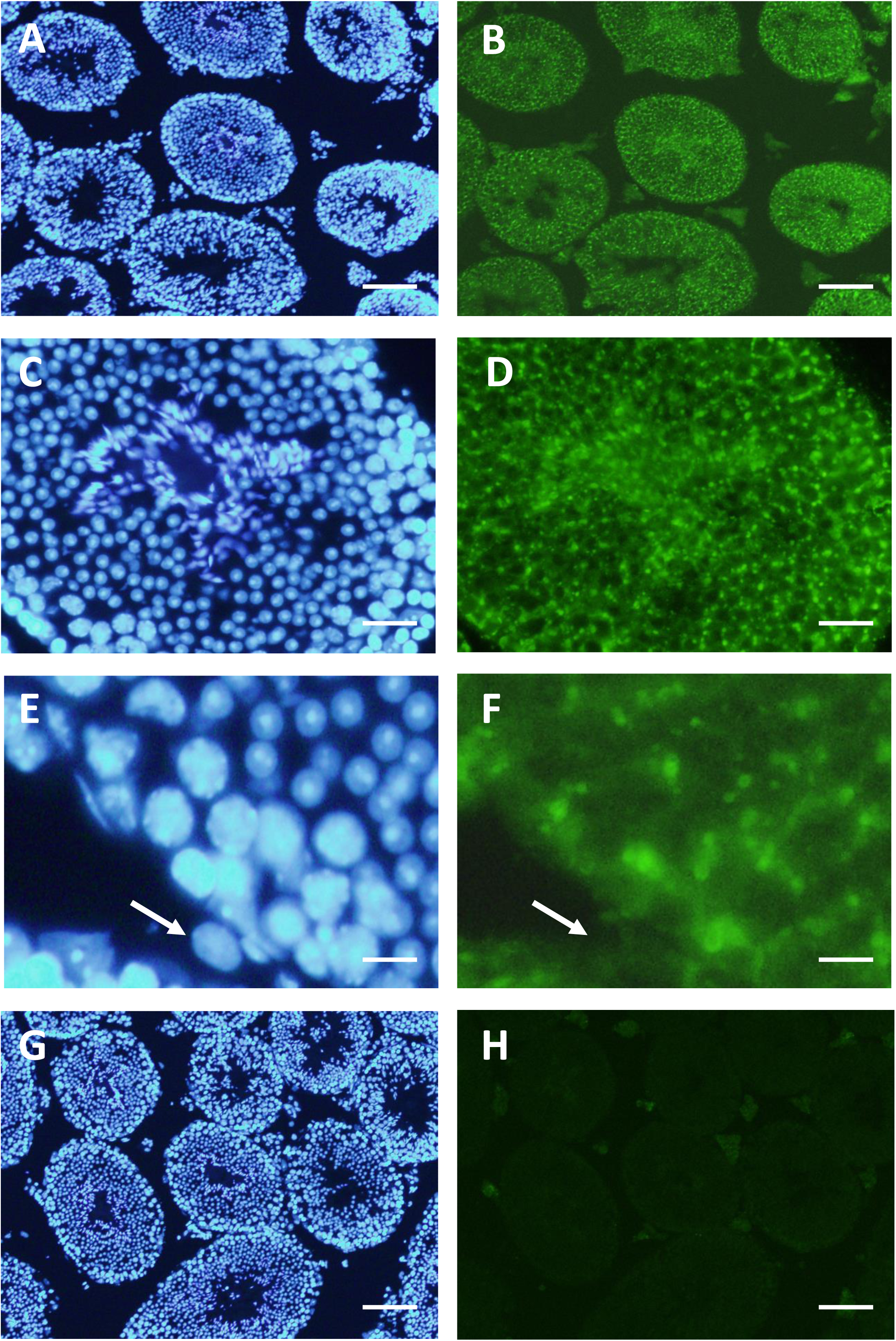
Immunohistochemistry of testis using anti-Lypd4 monoclonal antibody. In the testis, Lypd4 protein is found to localize to intracellular vesicles in germ cells at differentiation stages from spermatocyte to spermatid in the seminiferous tubules (B, D, F). Arrows in (E) and (F) indicate spermatogonia, which is Lypd4 negative. Such staining is not detected in the testes of KO mice at all (H). Control Hoechst staining are shown in A, C, E, and G. Magnifications, A, B, G, H, ×10; C, D, ×40; E, F, ×60.

**Supplemental Figure 5.**
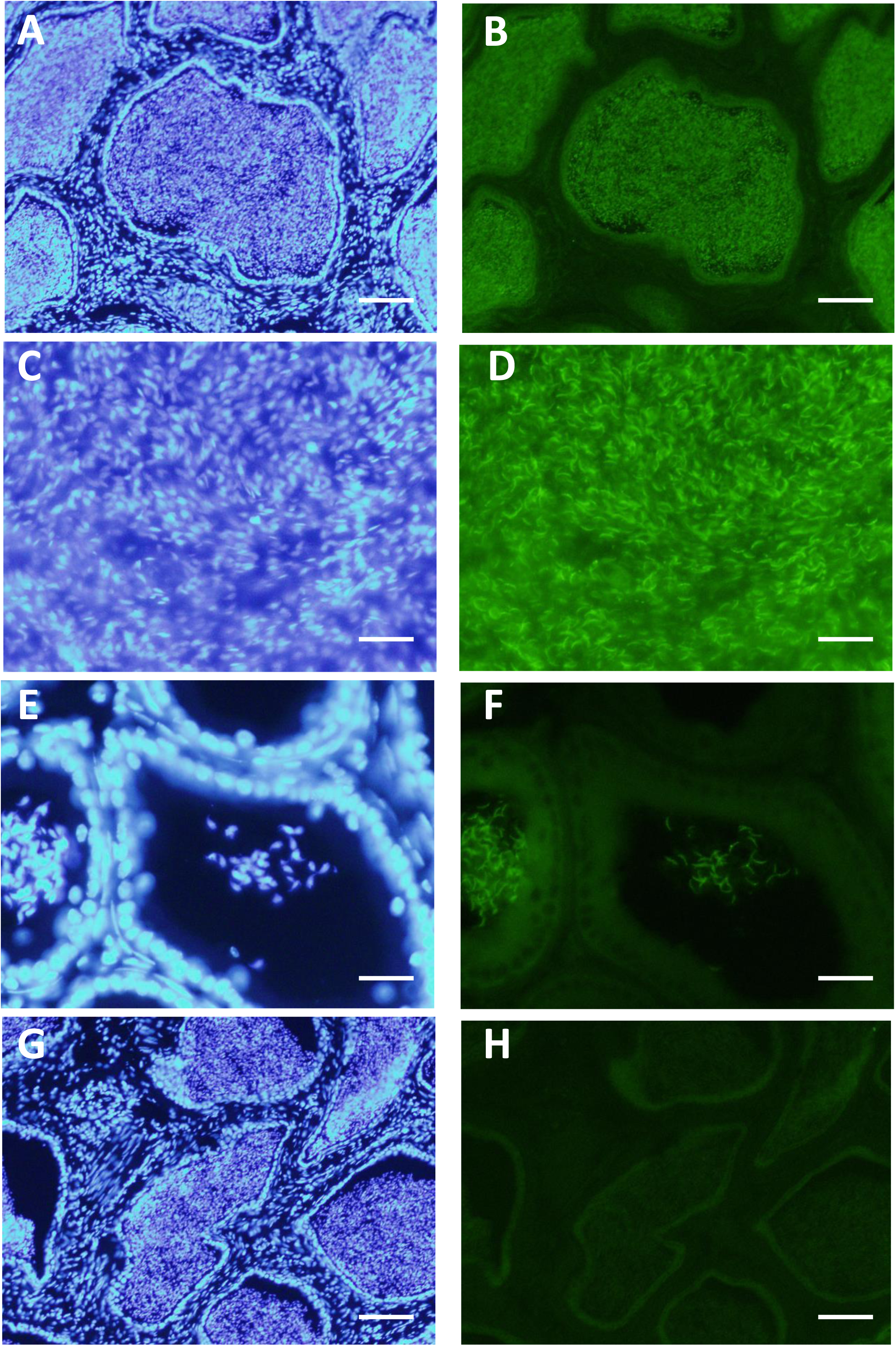
Immunohistochemistry of epididymis using anti-Lypd4 monoclonal antibody. In the epididymis, Lypd4 protein is found to localize to intratubular sperm (B, D, F). In the caput epididymis with low sperm number, the localization of Lypd4 to the sperm head is clearly visible (E, F). Such staining is not detected in the epididymis of KO mice at all (H). Control Hoechst staining are shown in A, C, E, and G. Magnifications, A, B, G, H, ×10; C, D, E, F, ×40.

**Supplemental Figure 6.**
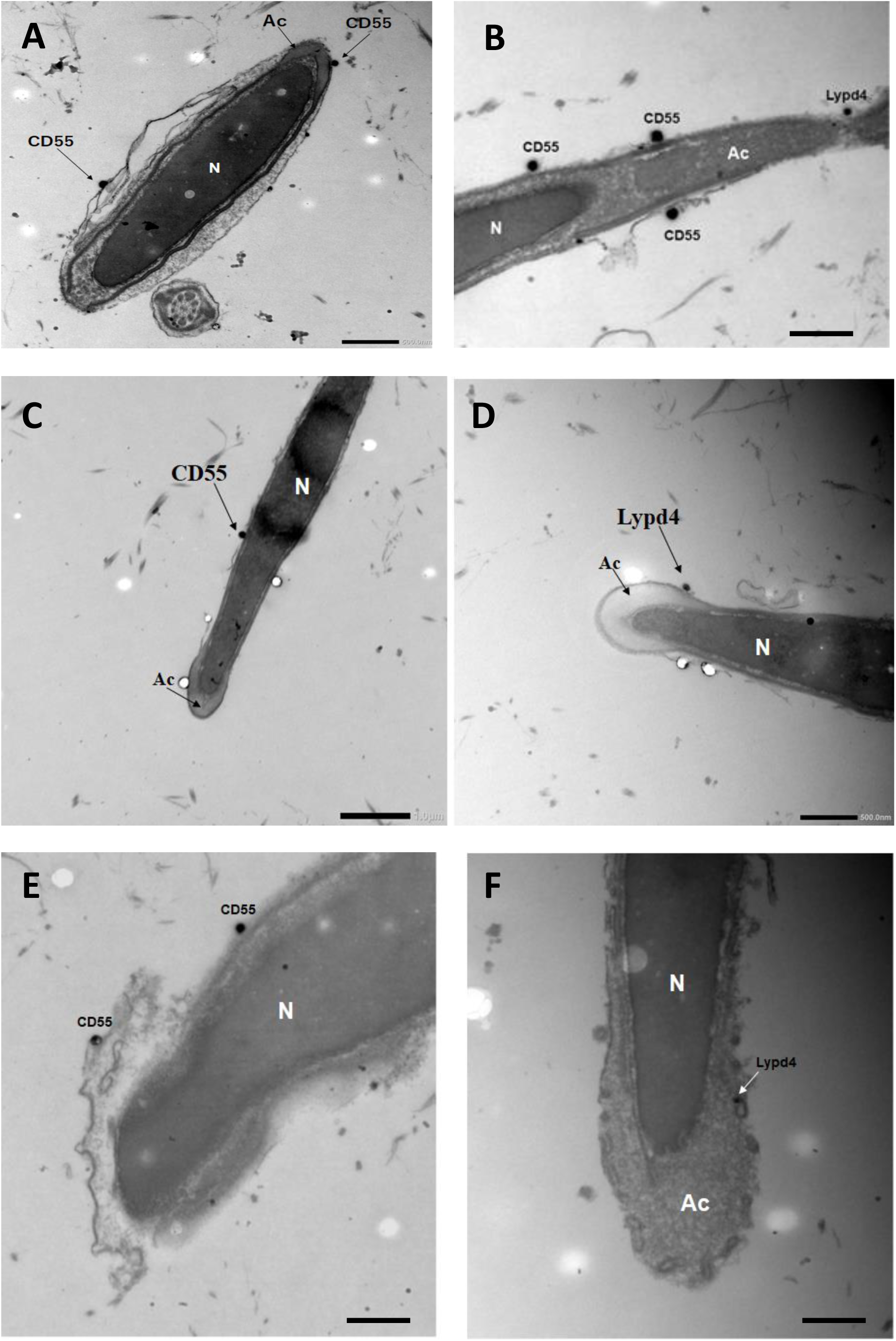
Immuno-electron microscopy of sperm using anti-Lypd4 and CD55 monoclonal antibodies. Extended data of Fig. 3L. Multiple acrosome-intact sperm (A, B, C, D) and acrosome-reacted sperm (E, F) were shown. Lypd4-positive or CD55-positive sperm are seen regardless of acrosome reaction. Scale bar, A, B, D, 500 nm; C, 1μm; E, F, 250 nm.

**Supplemental Figure 7.**
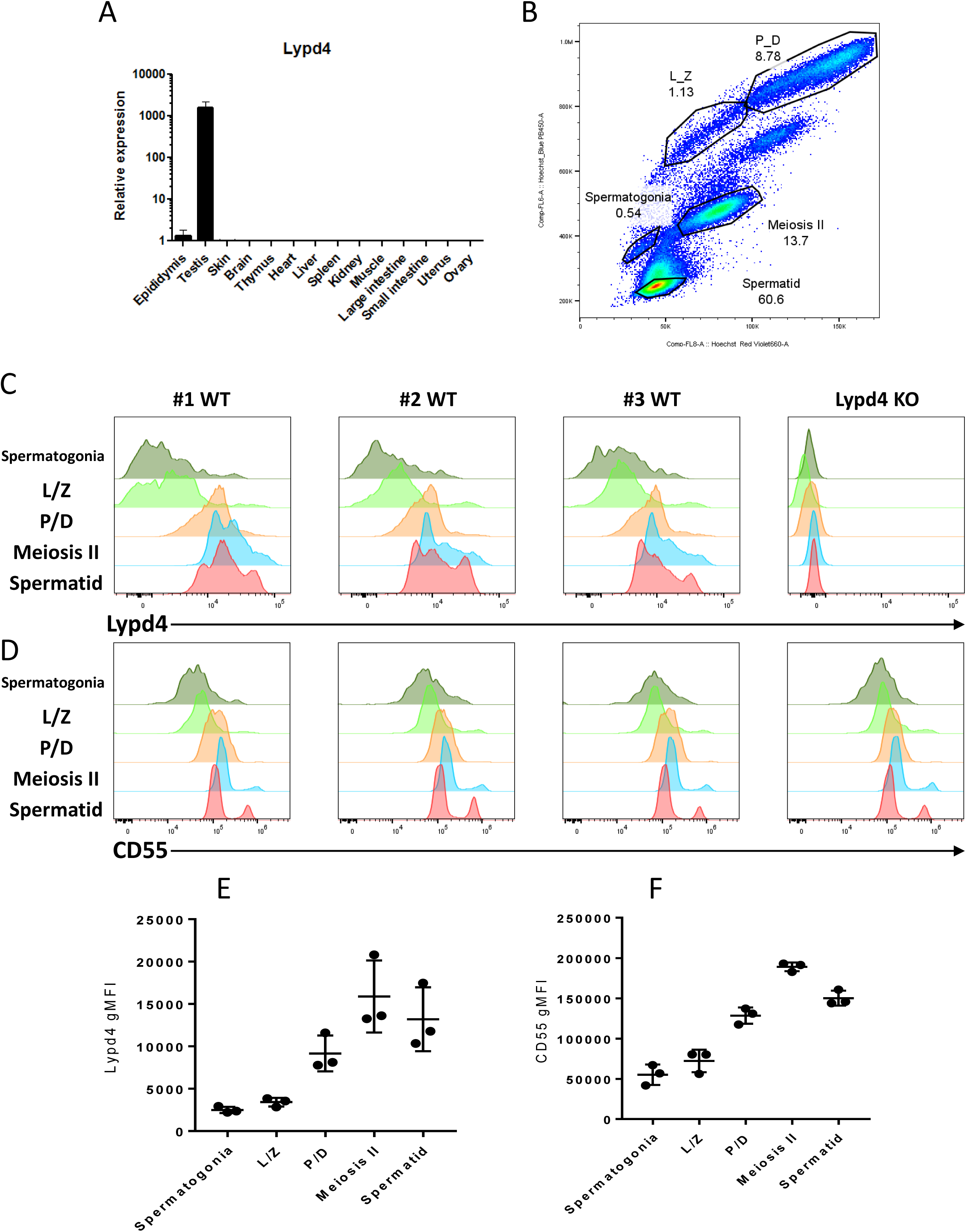
Expression of Lypd4 and CD55 in testicular cells. (A) Expression of Lypd4 mRNA in the whole body, analyzed by Taqman probe, showing the relative value of the expression in brain as 1.0. Specific and strong Lypd4 expression is observed in the testis. (B) Two-dimensional fractionation of testicular cells by Dual Hoechst staining^37, 38^. The fractions of testicular germ cells at each stage of differentiation and their proportion of the total are indicated. (C) FACS analysis of Lypd4 and CD55 in germ cells at each stage of differentiation. Results from three independent wild-type (#1∼#3 WT) and Lypd4 KO mice are shown; individual differences in Lypd4 expression in meiosisII spermatocyte and spermatid are observed, while CD55 expression is stable and constant. (D, E) Geometric mean fluorescence intensity (gMFI) of Lypd4 (D) and CD55 (E) in testicular cells; the expression intensity of Lypd4 is different in the three mice, while that of CD55 is nearly constant. L/Z, leptotene to zygotene spermatocytes; P/D, pachytene to diplotene spermatocytes.

**Supplemental Figure 8.**
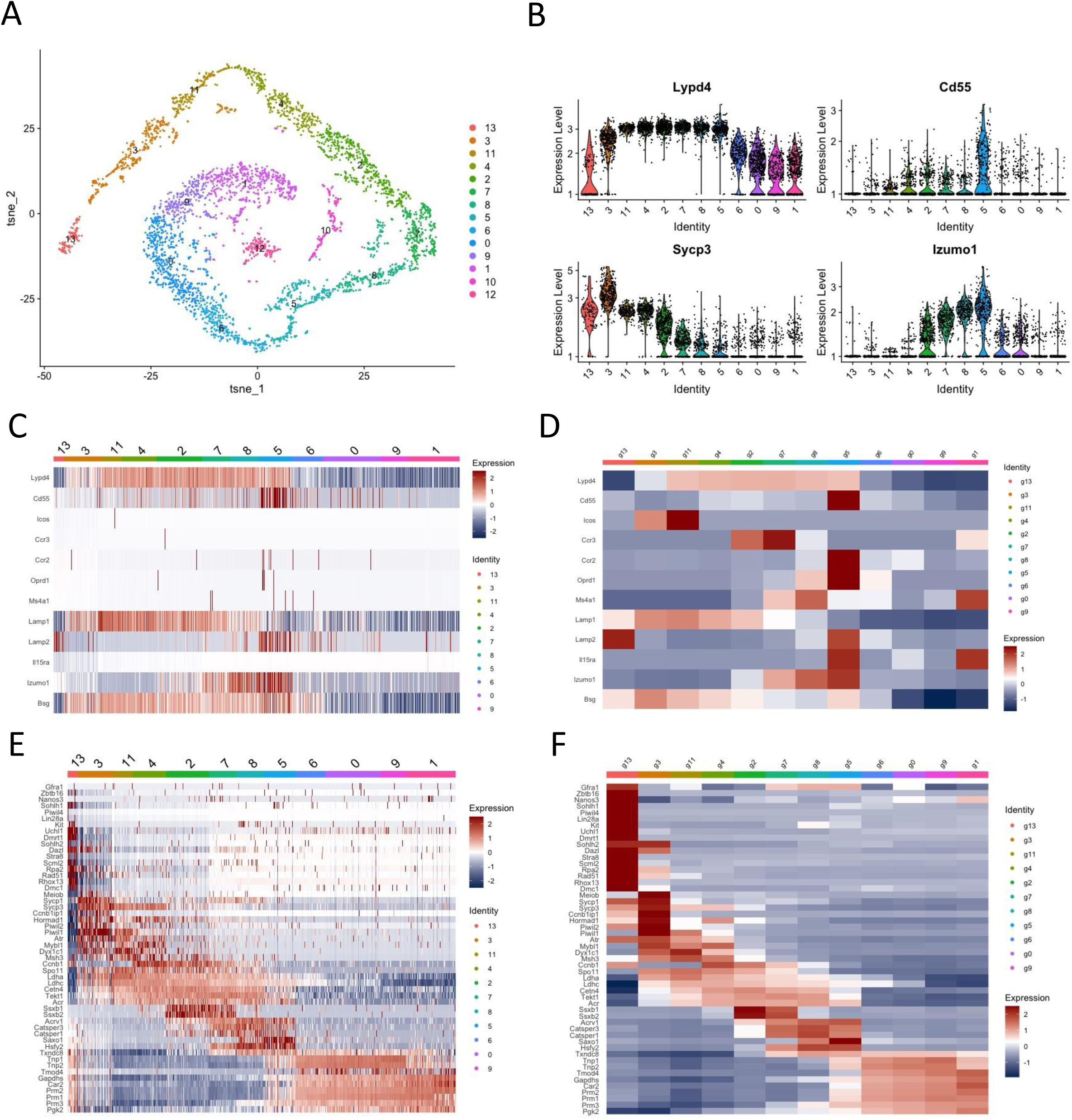
Single cell RNA sequencing re-analysis of mouse testicular germ cells. Lypd4 expression during mouse spermatogenesis was investigated by re-analyzing scRNA-seq datasets from GSM2928339, GSM2928340 and GSM292841^40^. (A) t-SNE plot illustrating the clustering of testicular cells across the datasets. (B) Violin plots depicting the expression levels of Lypd4 and indicated genes across the cell populations. (C, D) Heatmaps representing the expression profiles of Lypd4 and selected other genes across the cell populations. (E, F) Heatmaps showing the expression patterns of known spermatogenesis marker genes^40^. Cell clusters (13), (3, 11, 4) and (2, 7, 8, 5, 6, 0, 9, 1) correspond to spermatogonia, spermatocytes, and spermatids, respectively.

**Supplemental Figure 9.**
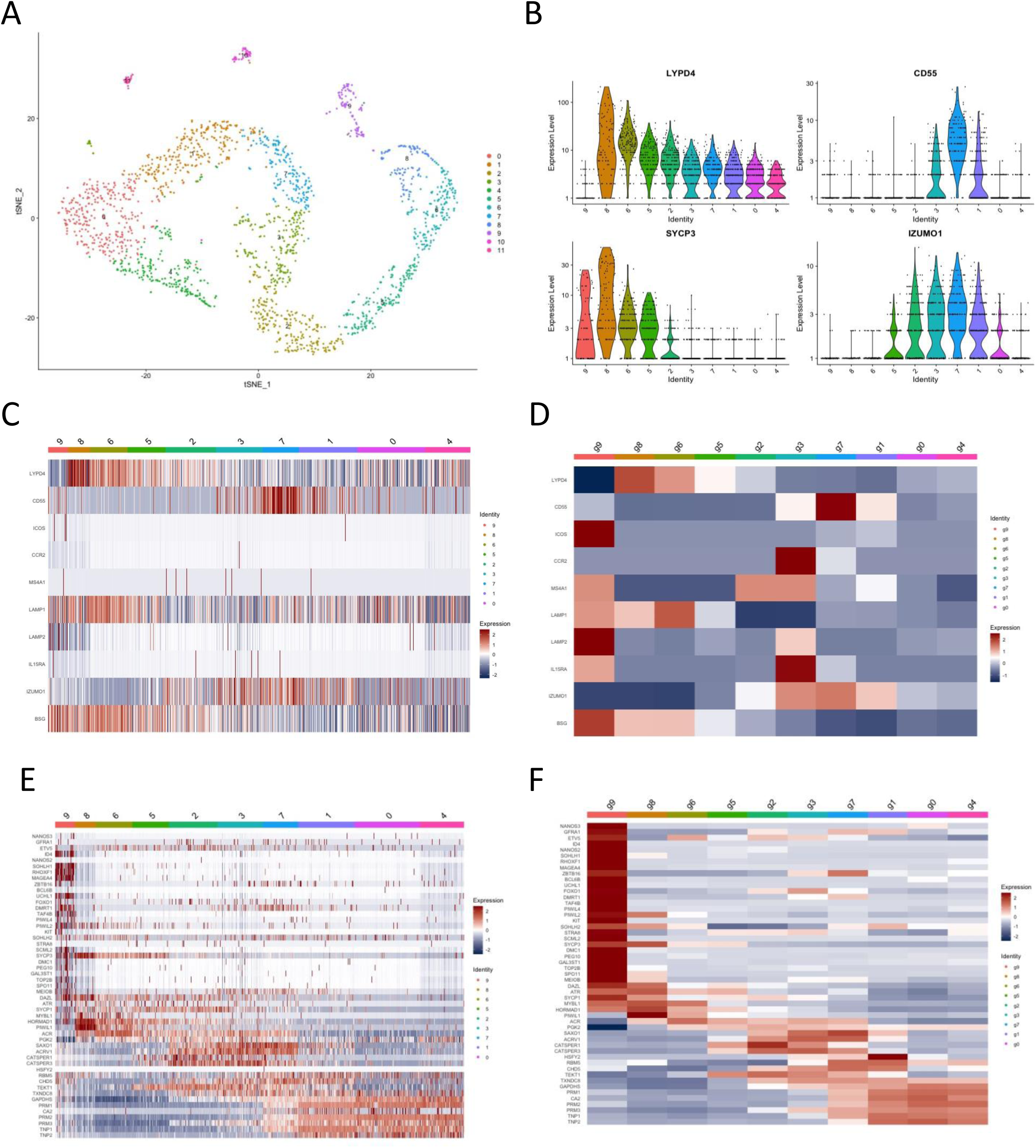
Single cell RNA sequencing re-analysis of human testicular germ cells. LYPD4 expression during human spermatogenesis was investigated by re-analyzing scRNA-seq data from GSM2928380^40^. (A) t-SNE plot illustrating the clustering of testicular cells. (B) Violin plots depicting the expression levels of LYPD4 and indicated genes across the cell populations. (C, D) Heatmaps representing the expression profiles of LYPD4 and selected other genes across the cell populations. (E)(F) Heatmaps showing the expression patterns of known spermatogenesis marker genes^40^. Cell clusters (9), (9,8,6,5) and (2, 3,7,1,0,4) correspond to spermatogonia, spermatocytes, and spermatids, respectively.

**Supplemental Figure 10.**
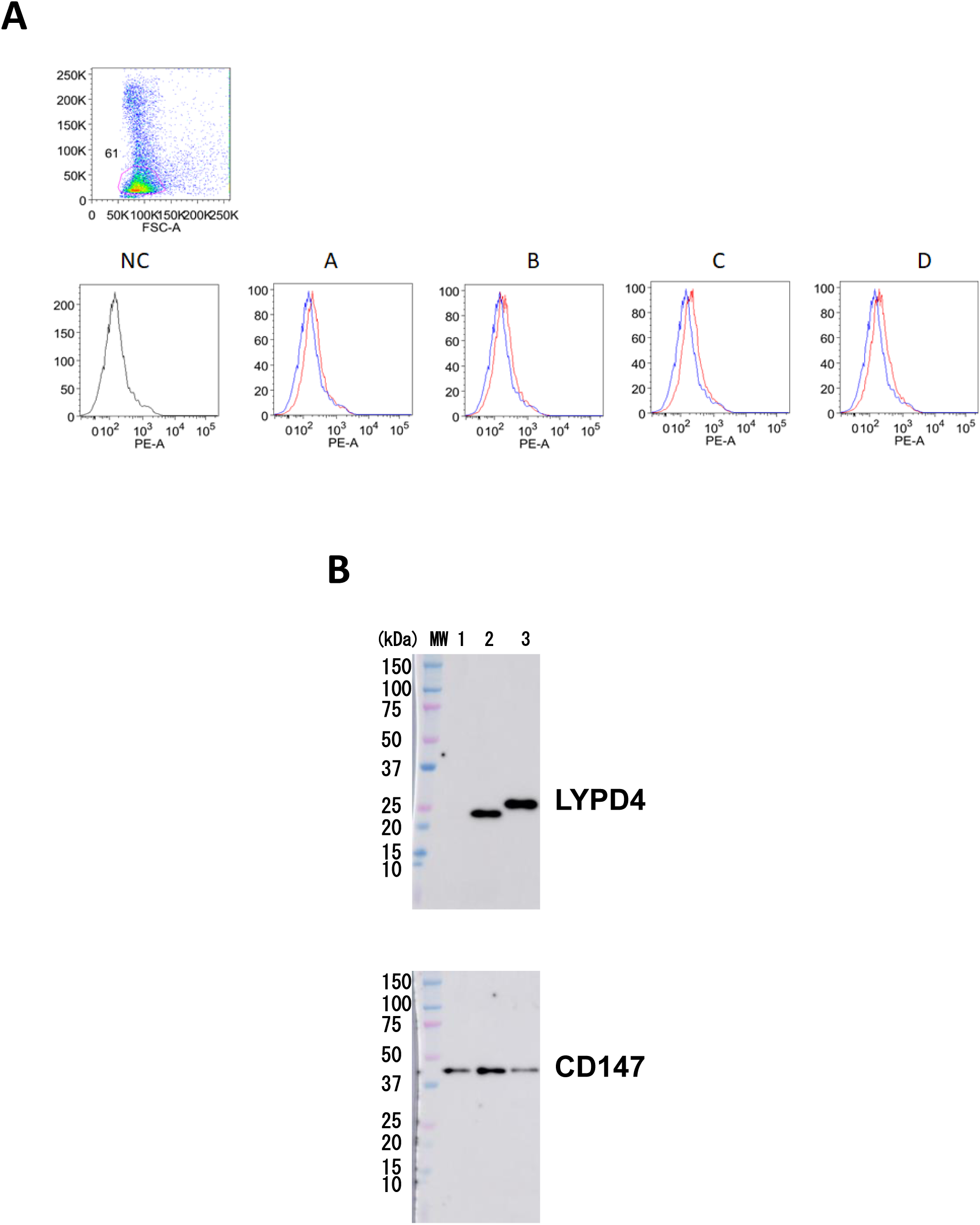
Generation of monoclonal antibodies against human LYPD4. (A) C57BL/6 mice immunized with Ba/F3-hLYPD4 cells were successfully immunized. Four independent clones of monoclonal antibodies were generated. However, all of them could not react with monkey sperm by FACS analysis. (B) Western blotting using clone A. It reacted with monkey testis (lane 2) and Ba/F3 cells expressing monkey LYPD4 (lane 3) but not with HTF-activated monkey sperm (lane 1). Quantitative controls by CD147 were displayed.

